# The Immunophenotype and Proviral Landscape of HIV-infected CD4 T Cells During Antiretroviral Therapy

**DOI:** 10.1101/2025.06.05.657210

**Authors:** Cyrille L. Delley, Sakshi Shah, Kevin M. Joslin, Yujung P. Park, Benjamin Demaree, Michael P. Busch, Mars Stone, Steven G. Deeks, Eli A. Boritz, Adam R. Abate, Iain C. Clark

## Abstract

In individuals on effective antiretroviral therapy, integrated HIV proviruses persist within CD4 T cells, forming a viral reservoir that rebounds if treatment is stopped. Identifying and targeting these rare, infected cells is critical for advancing therapies, but methods to study reservoir cells are limited and their unique properties remain largely unknown. We applied DAb-seq, a high-throughput method that combines single-cell DNA and surface protein sequencing, to profile over five hundred and twenty thousand CD4 T cells from the blood of six individuals on ART. Infected cells were unequally distributed in T cell subsets, and differential protein expression between infected and uninfected cells revealed significant heterogeneity across cell subsets. Attempts to identify surface markers that differentiate infected from uninfected cells found antigens that mirrored the enrichment of HIV in central memory subsets. However, while central memory T cells harbored the majority of HIV, cells with intact provirus were enriched relative to their defective counterparts in Naïve and Regulatory T cell subsets, suggesting that they differentially maintain intact proviruses. In summary, we developed DAb-seq as an open-source platform for linking the proviral landscape to diverse cellular phenotypes, revealing heterogeneity in surface protein expression and provirus maintenance across infected subsets.

## Introduction

Infection with human immunodeficiency virus (HIV) is incurable and remains a global health challenge. The HIV provirus reservoir is established rapidly after infection^1, 2^, persists in diverse tissues^3^, and is stable for decades^4–6^. Although antiretroviral therapy (ART) effectively suppresses virus replication, it does not eliminate the provirus reservoir, and lifelong treatment is required. While this fundamental obstacle to a cure is well established, the cellular mechanisms of provirus reservoir stability remain poorly understood. Insight into these mechanisms may inform therapeutic strategies to modulate infected cell proliferation, elicit HIV transcription for shock-and-kill, or promote infected cell detection and killing by immune cells. Methods to characterize the cellular state of infected cells are central to these goals.

Studying infected cells from people living with HIV (PWH) using functional or sequencing-based assays remains technically challenging. HIV+ cells in blood and tissue are rare (∼0.1-0.01% of CD4+ T cells)^7, 8^, and >95% of HIV+ cells harbor provirus with genomic defects and cannot produce infectious virions^9–11^. Unbiased isolation of HIV^+^ cells using fluorescence-activated cell sorting (FACS) is difficult due to an absence of unique cellular markers and the low expression of virus mRNA and protein during ART. Stimulation of infected cells can induce viral proteins for fluorescent labeling and FACS-based analysis, but activates only a subset of infected cells and alters their phenotype, preventing the study of cells in their natural state under ART^12–15^. High-throughput single-cell sequencing has shown potential for studying infected cells without stimulation, but challenges remain in identifying HIV^+^ cells in these datasets. Single-cell RNA-seq only detects infected cells expressing HIV RNA, and single-cell ATAC-seq only detects cells containing HIV DNA in accessible chromatin, limiting conclusions to a subset of infected cells. Neither of these measurements can accurately differentiate intact from defective provirus and therefore profile cells with mostly defective viruses^16, 17^. New methods are needed to study infected cells with intact provirus.

To address these challenges, we established an open-source high-throughput version of DAb-seq, a method we previously developed for single-cell DNA and protein sequencing on MissionBio’s commercially available Tapestri platform^18^. DAb-seq combines multiplexed PCR and DNA sequencing to link provirus intactness with the quantification of 153 surface proteins from the same cell, enabling detailed immunophenotyping of HIV-infected cells in their natural state. To increase the throughput of DAb-seq and analyze rare, infected cells at the single-cell level, we developed custom microfluidic devices and reagents that did not rely on a commercial instrument. We processed over 700,000 cells from six individuals living with HIV on ART, providing a more comprehensive analysis than previously possible. We found that provirus frequencies were highest in several memory T cell subsets and therefore implemented a statistical approach that accounted for the unequal distribution of infected cells across CD4 T cell subsets. Consistent with prior reports^17^, there was no evidence of specific surface expression patterns unique to infected cells. However, in contrast to a recent study^19^, we do not find signatures of immune selection at the surface protein level that are shared by cells with intact virus. Instead, proteins differentially expressed between HIV+ and HIV-cells were predominantly memory markers that reflected the unequal distribution of infected cells in memory subsets. Although infected central memory (CM) cells were the predominant reservoir of HIV provirus, they had a relative de-enrichment for intact provirus. Despite having fewer provirus overall, Naïve and regulatory T cell subsets showed relative enrichment for intact provirus. Our findings point to key differences in the fate of cells with intact proviruses between subsets and emphasize the importance of applying single-cell technologies that simultaneously measure provirus intactness and host cell phenotypes.

## Results

### High-Throughput Single-Cell Proteogenomic Analysis of HIV Reservoirs with DAb-seq

Under ART, HIV-infected CD4+ T cells are extremely rare, and greater than 95% of these cells harbor defective HIV genomes with large internal deletions, hypermutations, or single nucleotide polymorphisms (SNPs) that result in frameshifts or stop codons (**Fig. 1a**)^9, 15, 20^. Rarity, combined with the high prevalence of defective proviruses, complicates the identification of infected cells that cause viral rebound. To address this challenge, we single-cell sequence ten regions of the HIV provirus and surface proteins from the same cell (**Fig. 1b**). Barcoded primers are used to simultaneously amplify provirus genomic DNA and a 153-plex panel of barcoded antibodies, directly linking HIV genome sequences to surface protein profiles in single cells (**Fig. 1c,d**). This allows unbiased immunophenotyping of HIV+ cells in their natural state (**Fig. 1e**).

**Figure 1.**
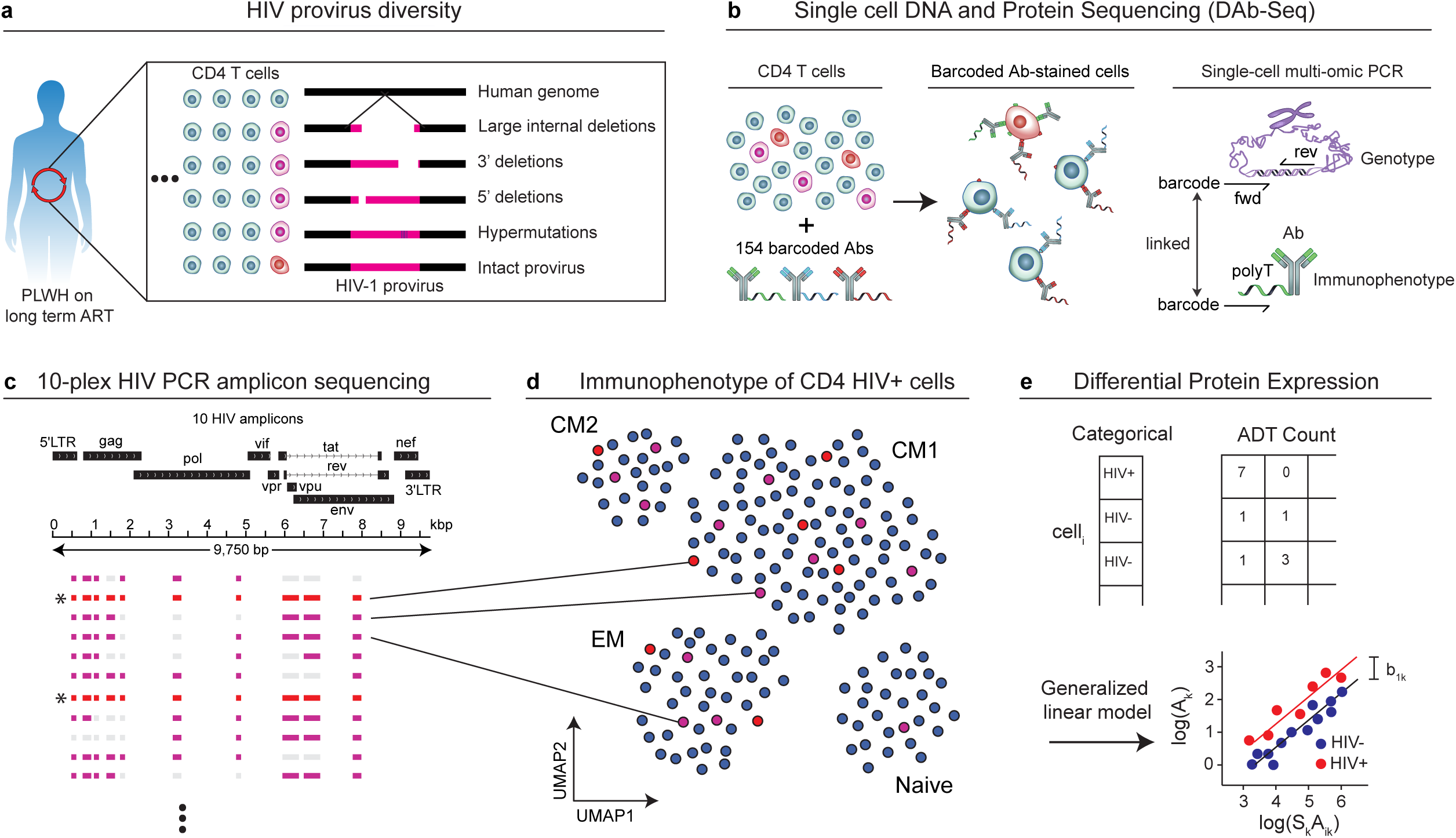
HIV DAb-seq workflow. **a)** Schematic depicting the diversity of HIV provirus sequences. **b)** Single cell DNA and Antibody Sequencing (DAb-Seq) sequences HIV and host cell PCR amplicons, and antibody derived tags (ADTs). **c)** Schematic showing the location and size of the 10 amplicons on the HIV genome. Intact and defective HIV genomes are inferred from the presence of HIV amplicons. **d)** HIV DAb-seq links rich single-cell immunophenotypes, cell infection status, and estimates of proviral intactness. **e)** Differential expression analysis between infected and non-infected cells aims to reveal HIV-specific surface antigen profiles.

The DAb-seq workflow employs a two-step microfluidic process to separate cell lysis and barcoding, providing the sensitivity needed to reliably amplify multiple regions of the HIV genome, a capability not achievable with common single-step methods (**Fig. 2**). First, cells are lysed with proteinase K to release genomic DNA inside a droplet (**Fig. 2a**). After heat inactivation of the protease, this cell lysate droplet is merged with a second droplet containing a barcoded bead with primers for DNA amplification (**Fig. 2b**). Barcoded primers are UV-released and amplify both genomic targets and antibody-derived tags (ADTs), linking these data in individual cells (**Fig. 2c**).

**Figure 2.**
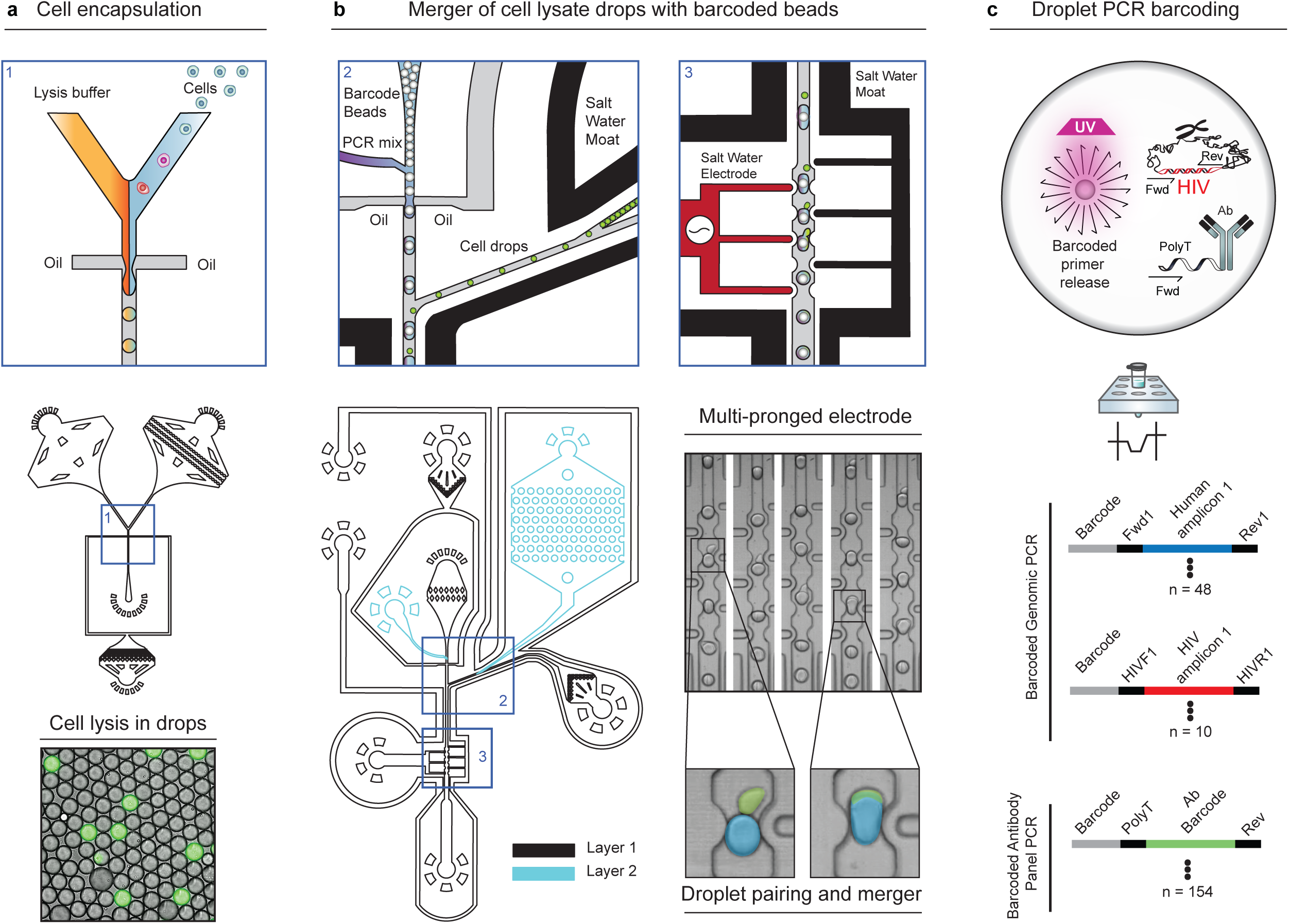
Custom microfluidic implementation of DAb-seq for HIV reservoir studies. **a)** Schematic of the cell encapsulation device (top) and SYBR-green stained droplets to confirm cell lysis (bottom). **b)** Schematic of the droplet merging device showing pairing of cell lysate drops with barcoded bead drops. **c)** Schematic of the barcoding droplet PCR reaction, including the barcode linkage between genomic PCR amplicons and antibody-derived tags.

We initially developed the DAb-seq workflow on Mission Bio’s Tapestri platform^18^. However, the commercial instrument has a typical recovery rate of around 20-40% of the cells loaded and requires repeated runs of 5-10k cells to reach sufficient throughput for large-scale reservoir studies, which is extremely costly. To address these limitations, we redesigned the workflow using custom microfluidic devices (**Supplementary Fig. 1**) and molecular reagents, achieving a tenfold increase in throughput that enables the processing of hundreds of thousands of cells per sample. The updated workflow features optimized microfluidics for cell encapsulation and lysis, and improved droplet merging with barcoded beads and PCR reagents (**Supplementary Data 1, 2**). We also incorporated enhanced barcode bead designs^21^ and unique molecular identifier (UMI) sequences in antibody-derived tags (ADTs). Collectively, these improvements increased data quality and significantly reduced costs. The new workflow achieves a median coverage of ∼3,300 UMI-corrected ADT counts per cell, compared to ∼750 uncorrected counts on the Tapestri platform^19^, which enhances the ability to detect differences in protein expression. This is crucial for correctly identifying cell signatures in HIV-infected cells. The system can process 100,000 cells in three hours for $7,774 ($0.08 per cell), compared to $61,755 (∼$0.62 per cell) using the commercial platform (**Supplementary Table 1**). Thus, our DAb-seq platform is an efficient and cost-effective solution for large-scale HIV reservoir studies.

### Validation of DAb-seq for Single Cell HIV Proteogenomics

To analyze the immunophenotype of HIV+ cells, we designed a multiplexed panel of HIV-specific primers targeting ten regions across the HIV genome, generating ∼300 base pair (bp) amplicons suitable for short-read sequencing (**Fig. 3a**). Our HIV amplicon panel covers 2,241 bps, representing approximately 24% of the HIV genome. Additionally, we included primers for 48 human genes to facilitate cell identification and normalize amplicon data (**Supplementary Table 2**). These human genes help distinguish cells from different donors and serve as internal controls.

**Figure 3.**
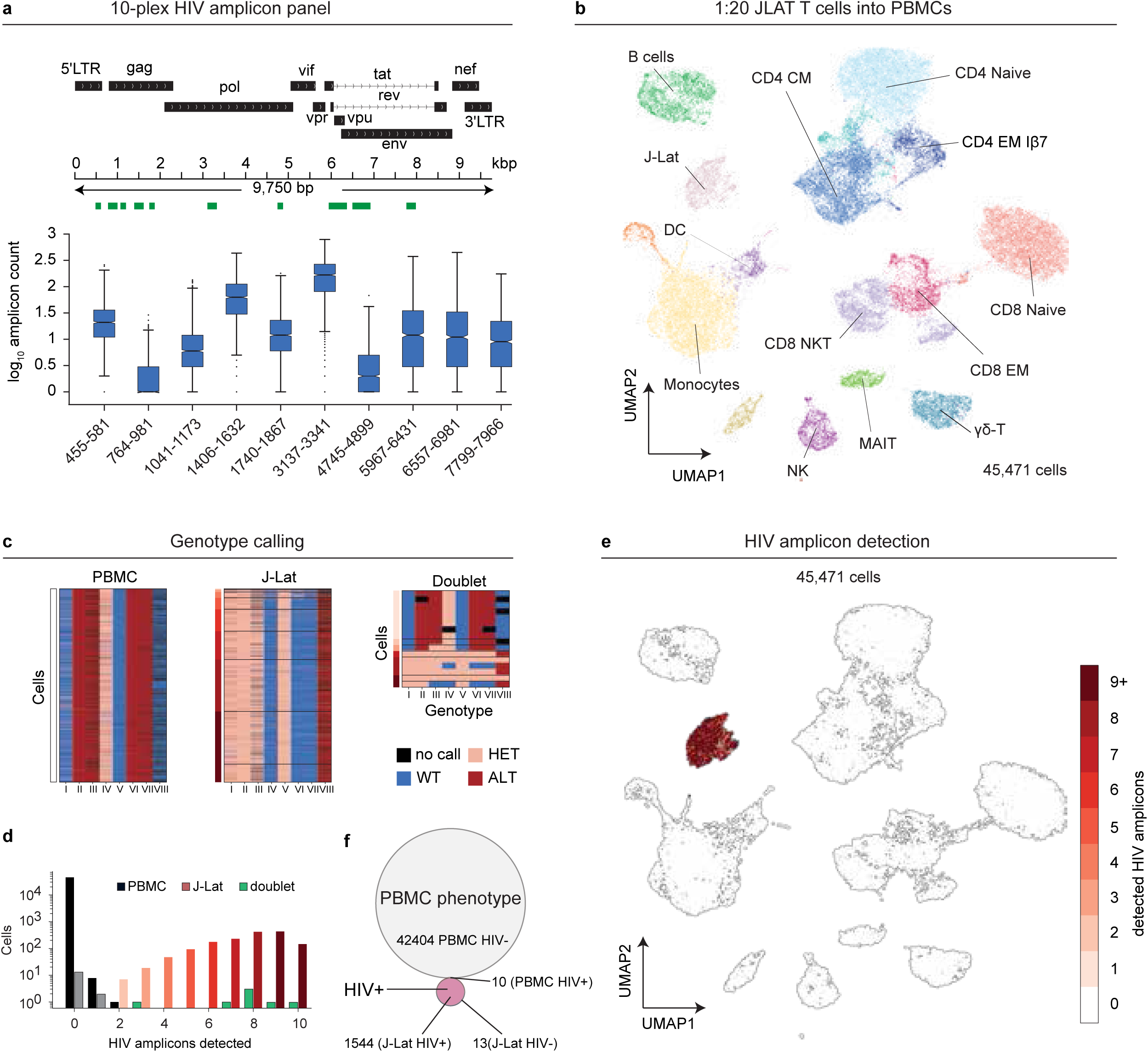
Workflow validation with a PBMC J-Lat spike-in experiment. **a)** Box plot of the number of sequenced amplicons per J-Lat cell. The notch indicates the median, the box is the inner quartile range (IQR), and whiskers are 1.5 IQR. Outliers are data points outside of these brackets. **b)** Uniform Manifold Approximation and Projection (UMAP) plot of J-Lat cells spiked into HIV-negative PBMCs. Colors indicate cell type. **c)** Heatmap showing variant calls (columns): wild type (WT), heterozygous (HET), alternate (ALT) or dropout for single-cells (rows). Mapped variants are, I: *IDH1* chr2-209113048 GA/G, II: *MOCS1-LINC00951* chr6-40116264 T/G, III: *SORCS3* chr10-32417945 G/A, IV: *WT1* chr11-32417945 T/C, V: *CES1P2* chr16-55770565 G/T, VI: *CES1P2* chr16-55770629 C/T, VII: *RAB31* chr18-9750662 T/C, VIII: *U2AF1* chr21-44524290 A/C. **d)** Bar chart depicting the number of sequenced HIV amplicons per cell. **e)** Identical UMAP as in b) with dots colored by number of detected HIV amplicons. **f)** Venn diagram showing overlap between J-Lat phenotype and detection of HIV genotype.

To characterize cell surface markers, we used a 153-antibody panel (BioLegend TotalSeqA), along with ten isotype controls to assess nonspecific binding, which enabled us to resolve the lineages of peripheral blood mononuclear cells (PBMCs) and determine cell states (**Supplementary Table 3**).

To validate our method, we spiked 2% HIV+ J-Lat cells into PBMCs from an HIV-negative donor, sequencing a total of 45,471 single cells. We developed custom bioinformatics scripts (**Data Availability**) to identify cell-specific barcodes and detect infected cells using HIV amplicon reads **(Supplementary Fig. 2a**). The ADT data were processed to identify feature barcodes and count UMIs, resulting in a raw ADT-by-cell count matrix. Major blood cell types, including CD8+ T cells, CD4+ T cells, natural killer (NK) cells, B cells, dendritic cells (DCs), and monocytes, were identified through marker expression and visualized via dimensionality reduction using Uniform Manifold Approximation and Projection (UMAP) (**Fig. 3b**).

J-Lat cells (1,544 cells) and PBMCs from the HIV-negative donor had unique genotypes that were identifiable by the 48 human gene amplicons, allowing us to distinguish between the two cell populations (**Fig. 3c, Supplementary Fig. 2b**). As expected, J-Lat cells exhibited a distinct immunophenotype characterized by high expression of CD4 and other T cell markers and were strongly enriched for HIV amplicons (median of eight amplicons per cell; 95% of J-Lat cells contained four to ten amplicons) (**Fig. 3d,e**). In contrast, only eight PBMCs contained HIV amplicons. However, PBMCs with more than two HIV amplicons displayed mixed single nucleotide polymorphism (SNP) profiles in the human gene amplicons, indicating likely co-encapsulation with a J-Lat cell (**Fig. 3f**). PBMCs with a single HIV amplicon matched the PBMC genotype, suggesting low background amplicon cross-contamination. Based on these observations, we set a threshold of two or more HIV amplicons to assign a cell as HIV positive. Using this criterion, we achieved a detection sensitivity of 99.17%, specificity of 99.98%, and accuracy of 99.95% for identifying HIV+ cells (**Supplementary Fig. 2c,d**). These metrics demonstrate the reliability of our improved DAb-seq workflow for detecting HIV genomes at the single-cell level.

### DAb-seq Analysis of HIV+ CD4 T Cells Under Effective ART

Next, we applied DAb-seq to study six people living with HIV (PLWH) who had maintained viral suppression (<50 RNA copies/mL) on ART for over two years (**Supplementary Table 4**). CD4 T-cells were isolated using negative selection from blood and sequenced using DAb-seq. ADT data were processed using CellBender^22^, and amplicon data were extracted and visualized following established protocols (**Supplementary Fig. 3**). After correcting for batch effects and integrating data across participants using totalVI^23–25^, we used seven distinct cell lines that had been spiked into each sample at 2% to confirm insignificant mixing between cell subsets (**Supplementary Fig. 4a,b**). In total, we analyzed 694,964 single cells with DAb-seq.

To determine cell subtypes, we performed unbiased clustering of surface markers using UMAP and subclassified CD4 cells using unsupervised community detection via the Leiden method^26^, label transfer from published single-cell blood datasets^27, 28^, and manual refinement (**Fig. 4a, Supplementary Fig. 4c-e**). Among the 526,755 CD4 T cells (56–88% of the total MACS-purified cells per participant), we identified 0.09–0.42% as HIV-positive per participant (1,182 in total) based on the presence of at least one HIV amplicon outside the long terminal repeat (LTR) region. Additionally, 977 cells were detected with only the LTR amplicon. Our control experiments using J-Lat cells had few LTR-only cells, supporting the conclusion that many infected cells in PLWH on ART contain only the LTR sequence. This is consistent with several recent reports documenting the prevalence of integrated solo LTRs in cells^29, 30^. HIV+ cells were found across all CD4 T cell subsets (**Fig. 4b-d, Supplementary Fig. 5**) with the highest enrichment in the central memory T cell subsets (CM1, CM2) and an effector memory subset expressing integrin-β7^31^. CD4 T cell Naïve, T reg, and cytotoxic (CTL) subsets contained fewer HIV+ cells, and only fourteen CD4^-^ HIV^+^ cells could be identified. Longitudinal samples from two participants showed a modest decline in infected cells over time that did not vary substantially between subsets (**Supplementary Fig. 6**).

**Figure 4.**
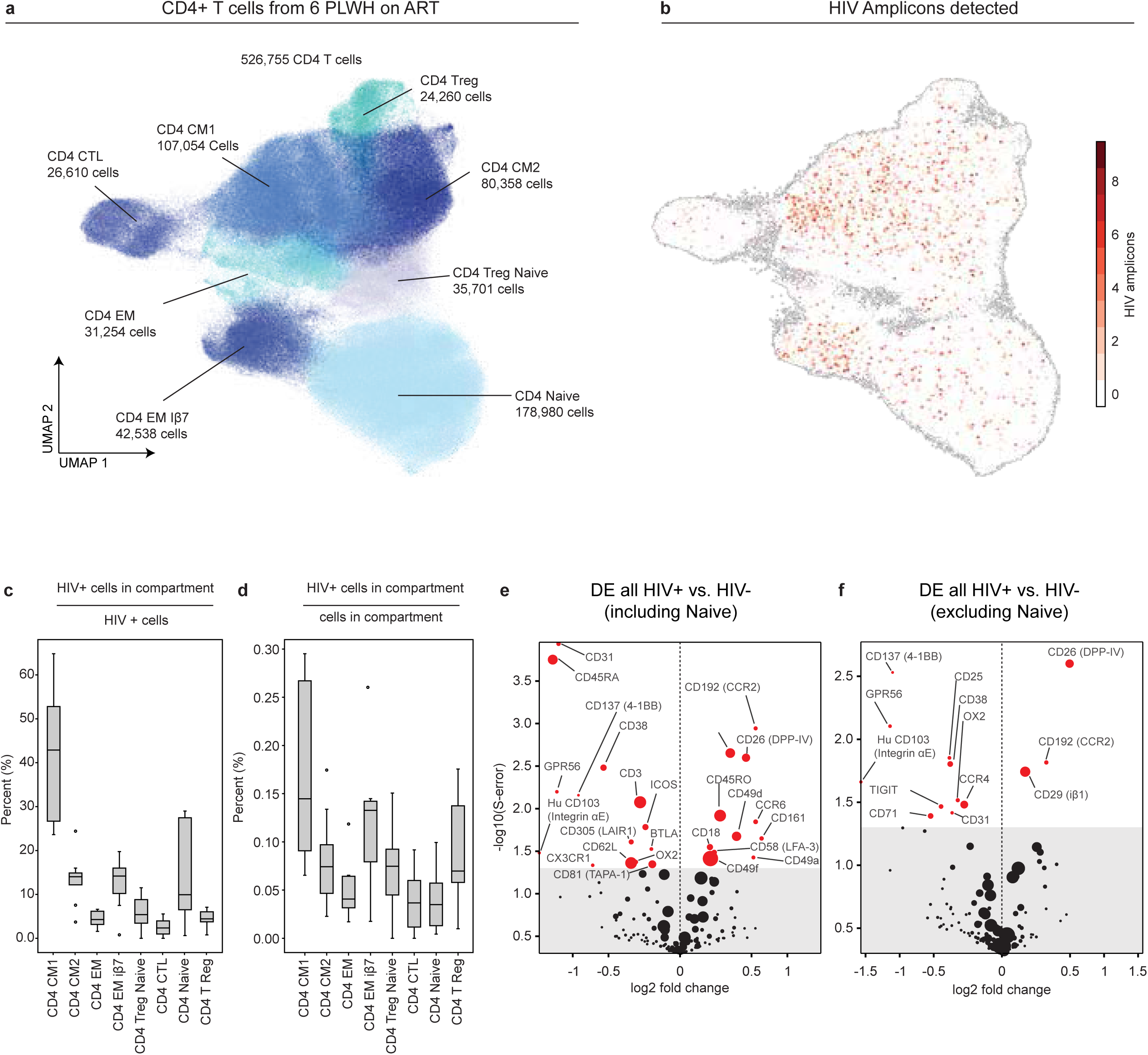
Frequency of HIV Infected Cells Within Different CD4 Cell Subsets. **a)** Integrated UMAP of the CD4+ T-cells from eight samples from six PLWH. Colors indicate CD4 subsets annotated with label transfer. **b)** Identical UMAP as in a) but colored with the number of detected HIV amplicons per cell. **c)** Percentage of HIV+ cells in each CD4 T cell subset versus all HIV+. **d)** Percentage of HIV+ cells in each CD4 T cell subset compare to the total number of cells in that subset. Boxes depict the mean and inner 5 to 95% quantile, the whiskers indicate inner 1 to 99% quantile. **e)** Differential protein expression analysis of all HIV-infected CD4 T cells versus all non-infected CD4 T cells. The volcano plot depicts the mean of the posterior estimate of the log fold change (x-axis) and the posterior probability of a sign error occurring. **f**) Same as in e) after removing CD4 Naive cells from the data set.

### Immunophenotypic Signatures of HIV+ CD4+ T Cells Under Effective ART

Discovering surface proteins that uniquely define all HIV+ cells would allow for the selective targeting of infected cells or their further characterization using functional and sequencing assays. DAb-seq identifies HIV+ cells by directly sequencing provirus DNA within the host cell, potentially enabling the detection of unique surface protein combinations that define these cells. However, previous studies have not yielded such markers^17, 19^, and HIV-infected cells in our data did not form a distinct group by unsupervised clustering (**Fig. 4b-d, Supplementary Fig. 4b**). Instead, infected cells were distributed across all major CD4+ T cell subtypes, indicating that infection status was not the dominant driver of clustering.

To search for linear or non-linear marker combinations that predicted infection, we partitioned data into training and validation sets and trained classification models (Neuronal-Net, random forest, Support Vector Machines) to predict which cell was infected based on antibody count data. When applied to the validation set, no classifier could predict HIV infection across all CD4 T cells or within individual CD4 subsets (**Supplementary Fig. 7**). Therefore, we conclude that no marker combination within the panel of 153 markers we analyzed can enrich for infected cells above the inherent enrichment of HIV observed in specific CD4 subsets.

Although we did not find a set of markers that predicted which cells were infected, we reasoned that differences in cell surface protein expression between HIV+ and HIV-cells might still exist. To search for these signatures, we performed differential expression analysis using a generalized linear mixed model. We treated ADT counts as a Poisson-distributed response variable and controlled for total UMI counts, donor identity, and infection status (**Supplementary Fig. 8a**) with linear predictors. This analysis revealed that HIV+ cells exhibited increased expression of 11 proteins (s-error < 0.05), including CD161 (KLRB1)^32, 33^, CD26 (DPP-4), the chemokine receptors CD192 (CCR2) and CD196 (CCR6)^34, 35^, and molecules involved in adhesion and trafficking, including CD58 (LFA-3) and the integrins CD49a (integrin α1), CD49d (integrin α4), CD49f (integrin α6), CD29 (integrin β1) (**Supplemental Table 5**). Additionally, infected cells had decreased expression of 14 proteins, including those with roles in modulating activation CD272 (BTLA), CD278 (ICOS), CD38, CD305 (LAIR1) and CD137 (4-1BB); and decreased expression of several proteins involved in T cell trafficking and adhesion, including CD62L (L-selectin), CD103 (integrin αE), CD31 (PECAM-1), and CX3CR1 (**Fig. 4e**).

### Controlling for HIV+ Cell Subset Distribution during Differential Expression

While our analysis identified proteins differentially expressed on HIV+ cells compared to HIV-cells, these signatures mirrored CD4 subset markers. Indeed, canonical markers for Memory (CD45RO) and Naive (CD45RA) CD4+ T cells were among the most differentially expressed proteins (**Fig. 4e**). Other markers known to be associated with CD4 T cell subtypes including Th17 (KLRB1, CCR6)^36^, Tissue Resident Memory cells (CD69, CD49a, and CD103), Central Memory cells (CD62L), and Effector Memory (CX3CR1) were also observed, suggesting that differential expression detected cell type associated proteins instead of true markers of infection. Notably, when Naïve cells were excluded from the analysis, many of these markers, including CD45RA and CD45RO, were no longer significant (**Fig. 4f**).

To understand if the unequal frequency of infected cells in CD4 T cell subsets was a major driver of differential expression results, we performed a “mock HIV+” analysis. For each HIV+ cell, we randomly selected a HIV-cell from the same subset, creating a “mock HIV+” group that contained no actual infected cells but matched the observed subset distribution of true HIV+ cells. We performed differential expression between the mock group and HIV-cells and observed a similar set of proteins to the previous analysis (**Fig. 5a,b**). The correlation between these analyses was high (R² = 0.83), suggesting that differential expression between HIV+ and HIV-cells resulted from the enrichment of HIV+ cells within specific subsets rather than a signature unique to HIV infection (**Fig. 5c**). Consistent with this, within-subset differential expression of HIV+ *vs* HIV-cells, and of HIV+ cells with intact virus *vs* HIV+ with defective virus (e.g. Naïve HIV+ vs Naïve HIV-) identified heterogeneous surface antigens that were not shared across all subsets (**Supplementary Fig. 9).**

**Figure 5.**
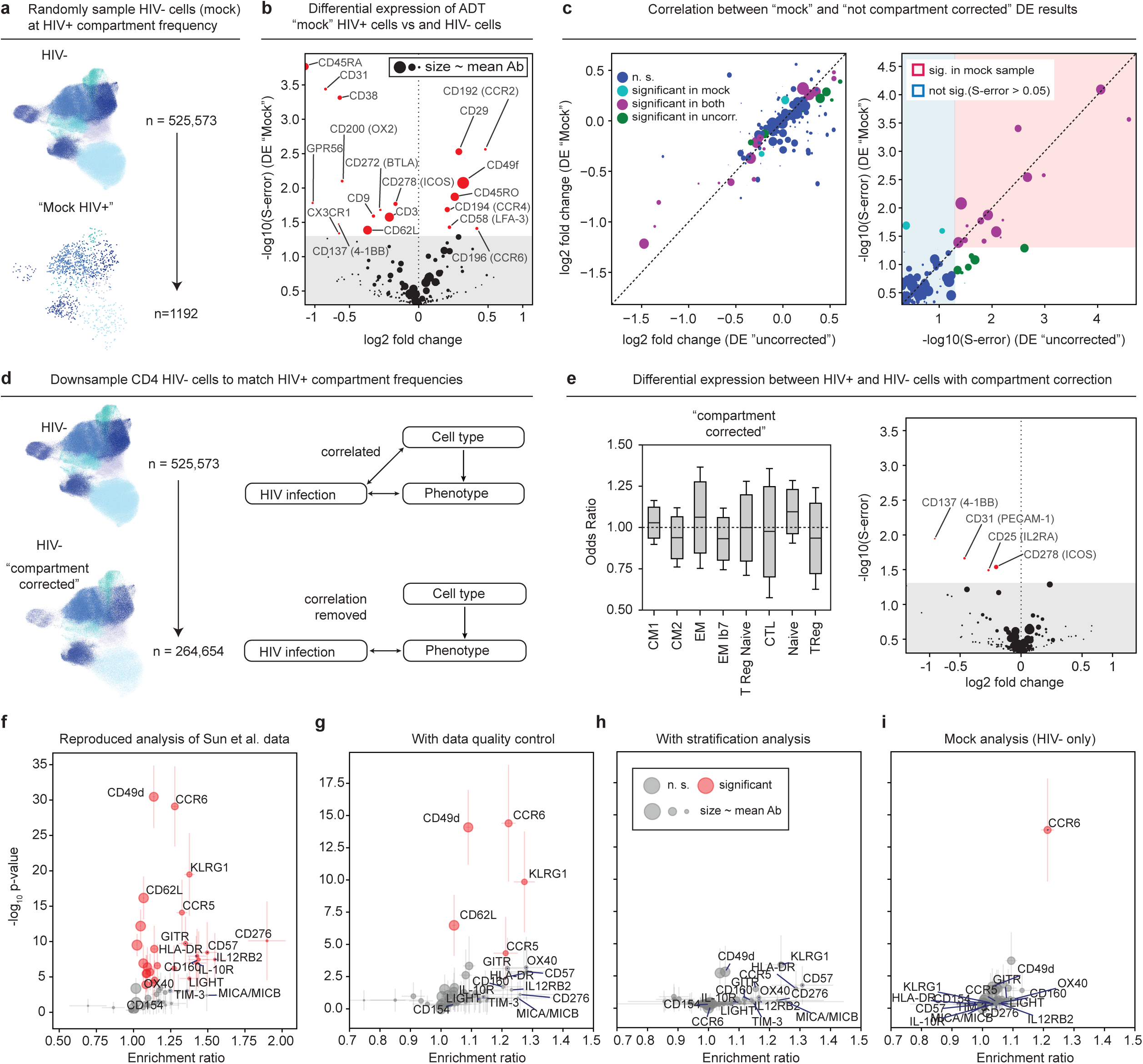
Controlling for the Effect of Infected Cell Subset Distribution during Differential Expression Analysis. a-c) Mock HIV analysis with only HIV-cells. HIV+ cells were replaced with HIV^-^ cells (mock) with the same cell subset distribution. Differential expression (DE) between mock cells and uninfected cells compared to the original DE analysis. **a)** Scheme used to randomly sample “mock” HIV+ cells (n=1192) from the HIV-cell population (n=525,573). **b)** Volcano plot of the differential expression analysis between “mock” HIV+ cells and HIV-cells. **c)** Correlation of the log2(fold change) and log10(S-error) between “mock” DE results and the original analysis. S-error has a similar meaning to p-value (see methods). **d,e) Stratified differential expression analysis.** The HIV-negative comparator was downsampled to match observed HIV+ subset frequency. **d)** Schematic used to select HIV-cells for the comparator group during differential expression analysis. **e)** Odds ratio of the HIV-comparator group after downsampling. The frequency of HIV+ and HIV-cells within each subset becomes approximately equal (Left). Volcano plot of the differential expression analysis between HIV+ cells and HIV-cells in the stratified analysis (Right). Compare to Figure Fig. 4e that contains no stratification. **f-i**) **Re-analysis of Sun et al. (2023) data. f)** Volcano plot replicating the analysis published in Sun et al. (2023) with their data and our implementation of their differential expression analysis method. **g)** Same analysis after removal of low-quality cells. **h)** Our analysis of Sun et al. data using stratification to remove cell type correlations. **i)** Our “mock” analysis of Sun et al data.

To define signatures of infected cells that were independent of cell subset type, we implemented a “stratification” approach. Instead of comparing all HIV+ cells to all HIV-cells, we subsampled HIV-cells to match the subset distribution of HIV+ cells (**Fig. 5d**). After applying this stratified analysis, only four proteins: CD25 (IL2RA), CD31 (PECAM-1), CD137 (4-1BB), and CD278 (ICOS) had significantly lower levels of expression in HIV+ cells compared to HIV-cells. However, the expression levels of these proteins were low, with less than twofold changes observed (**Supplementary Fig. 10**). The fact that most differentially expressed proteins were no longer significant after stratification supports the conclusion that differential expression between HIV+ and HIV-cells is dominated by differences in infection frequencies across cell types.

A recent study by Sun et al. (2023)^19^ employed DAb-seq with a different antibody and HIV amplicon panel on a commercial instrument to study the immunophenotype of HIV+ cells. That work identified many surface proteins as differentially expressed between HIV+ and HIV-cells with high statistical significance. Given the similarities between the studies, we reanalyzed their data, which contained 530,143 cells from five participants and 53 antibodies. Using their bioinformatic approach, we successfully replicated the reported findings (**Fig. 5f, Supplementary Fig. 11**). However, we noted their dataset had low antibody counts, high levels of isotype controls, and merged droplets containing multiple cells that required additional quality control measures (**Supplementary Fig. 12d-j**). We also note unequal ADT count distributions between intact, defective, and uninfected cells, which may impact statistical analysis (**Supplementary Fig. 12f,g**).

After applying standard quality control filters, such as thresholds for minimum antibody counts and exclusion of multiplets, we retained 360,117 cells (67.9% of the original dataset) for further analysis (**Supplementary Figs. 12**). In this adjusted dataset, we identified five proteins as statistically upregulated in HIV+ compared to HIV-cells: CCR6, CD49d, KLRG1, CD62L, and CCR5 (**Fig. 5g**). Notably, some of these markers also appeared in our study’s un-stratified analysis, suggesting potential cell subset effects. Therefore, we applied our stratification approach to their data, which adjusts the HIV-cell comparator to match the subset distribution of HIV^+^ cells and ensure equal representation of cell subsets in the analysis (**Supplemental Fig. 12k-m**). After this correction, no proteins remained significantly differentially expressed (**Fig. 5h, Supplemental Fig. 13**). Similarly, only CCR6 remained after “mock” analysis (**Fig. 5i**). We conclude that higher HIV+ cell frequencies within specific memory subsets strongly influences differential protein expression results.

### Immunophenotypic Signatures of Cells with Intact Provirus

The analysis of our data (**Fig. 4**) included cells with both intact and defective provirus, and results applied predominantly to cells harboring defective virus, which make up the vast majority of infected cells. We reasoned that this may have masked phenotypic differences that existed between cells harboring an intact provirus and cells with a defective or no provirus. To explore this, we classified cells based on the number of HIV amplicons, binning cells containing the top 10% of amplicons. Although some of the HIV genomes in these cells likely had genetic defects outside the detected amplicons, we refer to these cells for simplicity as having intact provirus. Next, we performed a differential expression analysis between cells with intact and defective proviruses. Initially, seven markers: CD9 (TSPAN 29), CD36, CD73 (Ecto-5’-nucleotidase), CD146 (MCAM), CD196 (CCR6), CD69 (type II C-lectin receptor), and CD272 (BTLA) were significantly downregulated on cells with intact proviruses compared to defective proviruses. After stratification to correct for cell subset effects (**Supplementary Fig. 14a-d)**, five markers (CD9, CD36, CD73, CD146, and CD196) showed significantly lower expression, while CD69 and CD272 were no longer significant (**Supplementary Fig. 15**). We also reanalyzed the Sun et al. (2023) dataset using their bioinformatic approach to compare the expression of proteins between cells with intact and defective provirus in their dataset. After correcting for subset distribution, this analysis yielded no significantly differentially expressed proteins (**Supplementary Fig. 16**).

### The Distribution of Intact Provirus within CD4 T cell subsets

Given the evidence that host differential expression results were driven by the enrichment of infected cells with CD4 subsets, we next asked if the relative abundance of infected cells varied with HIV genome “intactness” in each subset. We categorized and binned single cells based on the number of amplicons detected (LTR only, >0-30, 30-60, 60-90, >90 percentile) (**Fig. 6a,b**). In each bin, we computed the odds ratio, a measure of the relative likelihood of finding a cell with a given level of intactness in each CD4 T cell subset (**Fig. 6c**). Interestingly, the relative abundance of cells harboring intact viruses differed across cell types. For example, trends were observed within the CM2 subset, which had decreasing odds ratios with increasing levels of intactness, and within the Naïve, Naïve T reg, and T reg subsets, which had increasing odds ratios for increasing levels of intactness (**Fig. 6c**). Cells within the CM1 subset, which contained the highest number of HIV+ cells, had a relatively stable odds ratio across levels of intactness but a decrease in the odds ratio of the most intact virus (>90% amplicons). These trends held when we computed the odds ratio with respect to LTR-only cells, which provides a measure of the relative likelihood of finding a cell with a given level of genome intactness compared to infected cells with only LTR sequences **(Fig 6d)**. LTR-only cells are a valuable comparator because they track the natural cell dynamics of the infected cell population without the effects of provirus gene expression. Together, these results suggest that CD4 Naïve and T reg cell phenotypes, despite having far fewer total number of infected cells, may favor the persistence of more complete HIV genomes. We wondered if the processes driving this enrichment might be reflected at the surface proteome level and attempted to compare cells with intact provirus to cells with defective provirus within the Naïve and Treg subsets, However, the relatively small numbers of infected cells with intact genomes in these subsets (10 in T reg, 25 in Naïve) made this statistical analysis difficult. We also compared odds ratios for each HIV DNA amplicon individually (**Supplemental Fig. 17)** but did not observe striking relationships between specific amplicons and odds ratios for HIV+ *vs.* HIV- and HIV+ *vs.* LTR-only, with the exception of the amplicon targeting the matrix protein p17 (1041-1173), which was slightly reduced in CD4 EM, CD4 Treg Naïve, and increased in CD4 Treg. Taken together, these results suggest that the proviral landscape evolves differently in CD4 subsets. However, since the odds ratio compares the frequency of cells against the mean frequency, the observed relative enrichment of more intact HIV in CD4 Naïve and T reg cell phenotypes could result from the relative de-enrichment of intact HIV in other subsets, such as CM1.

**Figure 6.**
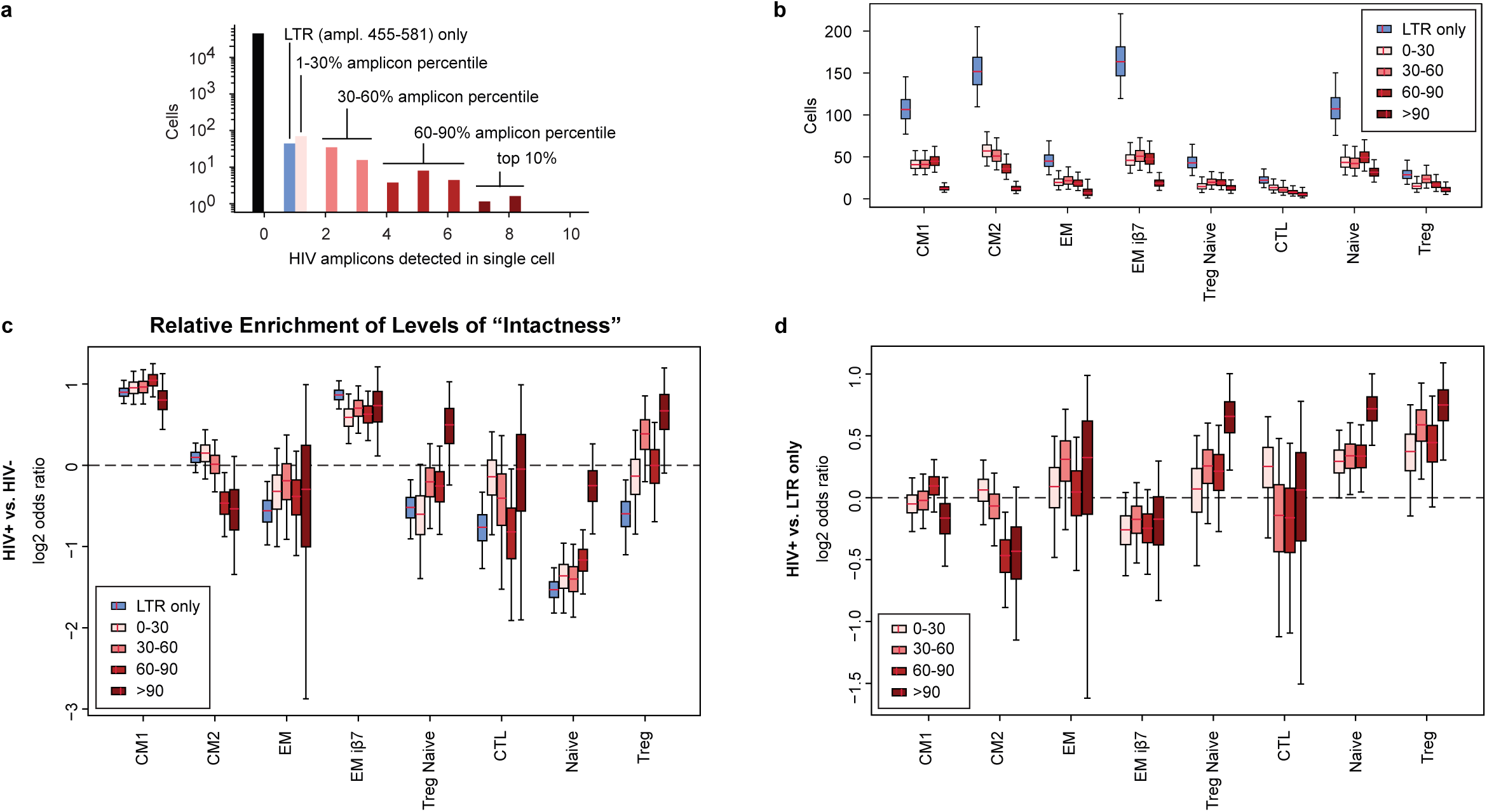
Differences in the Provirus Landscape of CD4 Subsets. **a)** Distribution in the number of HIV amplicons in single cells. Cells were binned by percentile into LTR-only, 1-30%, 30-60%, 60-90% and top 10% of HIV amplicons. **b)** Number of cells in each percentile bin for each CD4 T cell subset. **c,d)** Odds ratios of HIV genome “intactness” bins within CD4 subsets using HIV-(c) or LTR-only (d) cells as the comparator.

## Discussion

In the blood of PLWH on ART, HIV persists as an integrated provirus within CD4 T cells. Although this reservoir is believed to be a major source of virus rebound if ART is stopped, there is no consensus on the phenotype of infected CD4 T cells or the cellular determinants of persistence. Central to answering these questions is the ability to study single rare HIV+ cells from a background of diverse uninfected cells. This paper presents a powerful open-source technology for the high-throughput immunophenotype analysis of HIV DNA+ cells. Using this technology, we profiled 526,755 CD4 T cells from six individuals on long-term ART and precisely annotated CD4 T cell subtypes using label transfer techniques from a large single-cell CITE-seq dataset. HIV infected cells were phenotypically heterogeneous and distributed within all CD4 subsets but were enriched in a memory CD4 CM1 subset expressing integrin α4β1 (CD49d, CD29: VLA-4)^37^ and in a CD4 EM subset with high levels of integrin β7. The enrichment of HIV we observed within these cells may reflect the role of integrins in migration to sites of primary infection in inflamed tissues^38, 39^ and susceptibility to infection^40–44^, or natural T cell differentiation and turnover^45^. Longitudinal studies will be necessary to distinguish between these possibilities. In this context, DAb-seq is a powerful tool for future studies that track how the reservoir changes over time, in response to treatment interventions, or in individuals that control rebound.

We attempted to identify surface markers that were differentially expressed in all HIV+ cells, even despite the large immunophenotypic heterogeneity observed. Identifying protein markers of the HIV reservoir remains an important goal because it would facilitate the isolation of HIV+ cells for mechanistic studies and the design of therapeutic strategies to eradicate HIV. Previous work has shown modest associations between infection and immune checkpoint molecules (LAG-3, TIGIT, CTLA-4, PD-1)^46, 47^, activation markers (CD69, CD2, OX40)^48–50^, and several chemokine/cytokine receptors (CXCR5, CCR6) ^34, 51^, but no evidence that any protein, or combination of proteins, can uniquely identify HIV+ cells. We reasoned that the large number of surface proteins profiled in our study and the ability to accurately identify cells with integrated provirus might provide an advantage over previous studies. We performed differential protein expression analysis (DE) between all CD4 HIV+ and HIV-cells (with and without Naïve CD4 T cells) using a generalized linear model that accounted for variability in count data. This analysis found canonical markers of CD4 memory cell subsets, suggesting that the unequal distribution of HIV+ cells within memory subsets contributed to DE results. We verified this using two independent analyses. First, we replaced HIV+ cells in our analysis with HIV-cells sampled to match the number and location of infected cells within CD4 subsets (mock analysis) and found that, despite no HIV+ cells present in the data, DE results correlated strongly with the initial analysis. Second, we downsampled uninfected cells to create an HIV-comparator that matched the distribution of HIV+ cells within CD4 subsets (stratified analysis) and found that most differentially expressed proteins were no longer significant. Our analysis of the dataset from Sun et. al (2023) reached identical conclusions. These results strongly suggest that DE analyses detected proteins associated with the CD4 subsets enriched in proviruses, instead of signatures unique to infection and independent of cell type. This does not preclude the possibility such markers exist and can be captured using larger protein panels or epigenetic, RNA, or metabolic measurements. However, the information in genome-wide transcriptome and epigenome measurements will be equally confounded by the underlying HIV distribution within subsets. In this regard, our statistical approach provides a framework for separating these effects that can be applied to single-cell omics datasets.

A significant goal of single-cell studies is understanding cellular mechanisms contributing to persistence. Therefore, the rich phenotypes measured by single-cell technologies must also determine which cells have intact provirus. This is not trivial; commercially available scRNA-seq and scATAC-seq methods currently cannot sequence enough of the HIV genome to differentiate intact from defective virus. DAb-seq has the unique ability to barcode and sequence HIV PCR amplicons. Like IPDA, this information is a powerful, albeit imperfect, measure of HIV genome intactness. In our analysis, we binned infected cells based on the number of HIV amplicons detected and compared cells with more intact genomes (top 10%) to those with less intact genomes. We found markers (CD9, CD36, CD73, CD146, and CD196) that had statistically lower expression in cells with intact proviruses after correcting for cell subset effects. Although these proteins have potentially interesting connections to persistence, including regulating immune activation/exhaustion (CD36 and CD73)^52^, virus membrane budding and fusion (CD9)^53–55^, and adhesion/migration (CD146, CD196), they were weakly or infrequently expressed in our data, warranting cautious interpretation. This may be because, despite single cell sequencing over 525,000 CD4 T cells, we annotated only 125 as likely to contain intact HIV genomes. This highlights the difficulty in sequencing enough cells with intact provirus to detect statistical differences. Sorting methods, like the one we recently developed^56, 57^, aim to solve this problem by providing experimental access to more cells with intact provirus.

The forces that control reservoir persistence are poorly understood^45^. We reasoned that more stable reservoir cells would differentially harbor intact provirus and looked for cell types with relative enrichment in intact HIV genomes relative to defective ones. After binning cells by their level of HIV genome intactness and calculating the odds ratio, we noted that the Treg, Treg Naïve, and Naïve subsets had increasing odds ratios with increasing HIV intactness levels, suggesting that despite their relatively small reservoirs, these subsets favored the maintenance of more intact proviruses. Conversely, memory cells had the largest percentage of HIV proviruses but appeared less conducive to harboring intact provirus. This was especially pronounced in the CM2 subset. Previous work has shown that intact HIV proviruses decay faster than defective ones^7^, and that immune selection or cytotoxicity from HIV expression can affect decay ^29, 58^. Our results imply that this depends on the T cell subtype, with Naïve, Naïve T reg, and T regs being potentially more favorable for maintenance. These findings are consistent with reports that infected Naïve CD4 T cells produce comparable amounts of infectious virus to memory CD4 T cells ^59^ and add to the growing body of literature supporting the importance of Naïve CD4 cells in persistence^60–63^.

Our results have important implications for studies that define host persistence mechanisms, including DAb-seq, scRNA-seq and scATAC-seq studies. It is increasingly clear that the HIV reservoir resides non-randomly within phenotypically diverse CD4 subsets in blood and tissue^17, 19, 37^. Host signatures that reflect this distribution or that are specific to cell subtypes are interesting if conserved across people and may provide insight into preferential infection, differentiation, proliferation, and immune selection^45, 64^. However, a major goal of single-cell studies should be to discover mechanisms that can be targeted to eliminate reservoir cells within all subsets. Such mechanisms, if they exist, will be found by controlling for cell type and identifying host features dependent on intact virus.

## Methods

### Fabrication of microfluidic devices

Microfluidic devices (**Supplementary Fig. 1**, **Supplementary Data 1**,**2**) were fabricated with standard photolithography techniques at UCSF and the Biomolecular Nanotechnology Center cleanroom facility at UC Berkeley. Three inch or four-inch silicon wafers (University Wafer) were spin-coated (Laurell H6-23) *with* SU-8 2025/2050 photoresist (Kayaku Advanced Materials) and features were patterned using a UV mask aligner (OAI 206) and photomask transparencies (CAD/Art Services). After UV exposure, wafers were developed and baked overnight at 65°C. PDMS pre-polymer and curing agent (Dow Sylgard 184) were mixed in a 10:1 ratio, degassed, poured over the patterned silicon master molds, and degassed again to remove bubbles. PDMS devices were baked overnight at 65°C, cut out from the mold, and inlets/outlets were punched using a 0.75 mm biopsy punch (World Precision Instruments 504529). PDMS devices were bonded to 75 mm x 50 mm x 1 mm glass slides (Fisherbrand 12-550C) using an oxygen plasma system (Tergeo; PIE Scientific). After baking at 150°C for 5 min and a subsequent 65°C bake overnight, bonded microfluidic devices were treated with Aquapel (PGW Auto Glass) to make droplet-contacting channels hydrophobic.

### Preparation of barcode beads

Barcode bead fabrication has been described in detail^21^, with the following changes. The uracil containing release primer was replaced with the UV cleavable HDAB 39; ligation junctions of the barcode blocks have been replaced by optimized sets^65^. Final primer ligation was performed at 0.6 µM HIV primers (HDAB 45 – HDAB 54, 0.06 µM each), 1.2 µM AMLv2 fwd primers (0.025 µM each), 0.15 µM HDAB 42 primer for the antibodies and using HDAB 41 as splint at a concentration of 1.94 uM. All oligonucleotide barcode and primer sequences can be found in **Supplementary Table 2**.

### Human cell line culture

A 7-cell spike-in control was prepared by pooling 2.5 million cells from cultures of Jurkat, K-562, HuT 78, Raji, THP-1, U-937, KE37 cells. Aliquots were cryopreserved in cryopreserved in 0.5 ml of 90% FBS + 10% DMSO. Cell lines used in these experiments were Jurkat (ATCC 88042803), K-562 (ATCC CCL-243), HuT 78 (ATCC TIB-161), Raji (ATCC CCL-86), THP-1 (ATCC TIB-202), U-937 (ATCC CRL-1593.2), KE-37 (DSMZ ACC 46), and J-Lat Full Length Cells (6.3, ARP-9486; HIV Reagent Program). Jurkat, K-562, Raji, U-937, and KE37 and J-Lat cell lines were cultured in Gibco RPMI Medium 1640 with 10% VWR Fetal Bovine Serum, 100 U/mL penicillin G, and 100 μg/mL streptomycin (Gibco 15140122). HuT 78 cells were cultured in Gibco Iscove’s Modified Dulbecco’s Medium (IMDM) with 20% FBS and 100U/mL penicillin G and 100 μg/mL streptomycin. THP-1 cells were cultured in Gibco RPMI Medium 1540 with 10% FBS, 1% sodium pyruvate (NaPyr), 1% HEPES, 4.5g/L glucose, and 0.05 mM 2-Mercaptoethanol. Jurkat, K-562, HuT 78, Raji, THP-1, U-937, and KE37 cells were purchased from the Barker Cell Culture Facility at UC Berkeley.

### Cell preparation for human immune cell/J-Lat validation experiments

2 million peripheral blood mononuclear cells from a healthy donor were mixed with 0.04 million J-Lat cells and resuspended in 160 μL MACS Buffer in a Protein Lo-Bind Tube (Eppendorf #0030108116).

### Study Participants

Peripheral blood mononuclear cells (PMBC) from study participants were provided by the HIV Reservoir Assay Validation and Evaluation Network (RAVEN). Samples were collected from ART-suppressed people living with HIV under IRB-approved protocols, and all participants provided informed consent. Detailed clinical information about each study participant can be found in **Supplementary Table 4**.

### Handling of PBMC samples from study participants

Cryopreserved samples containing 50 x 10^6^ cells were shipped in liquid nitrogen to UC Berkeley. Samples were thawed at 37°C until only a small amount of ice remained and pipetted into 25 mL prewarmed 37°C RPMI Medium 1640 (Gibco 11875085) with 10% Fetal Bovine Serum (FBS; VWR S181B-500) and gently mixed by inversion. Cells were centrifuged for 10 min at 300 x g (Sorvall ST4R Plus-MD; Thermo Scientific 75009525) at 4°C, supernatant was aspirated, and cells were washed once in in MACS Buffer containing 2 mM EDTA (Sigma-Aldrich E4884-100G), 0.5% w/v BSA (NEB #B9200) in Dulbecco’s Phosphate Buffered Saline Solution (D-PBS, no Mg, no Ca) (Gibco 10010-023). CD4 T cells were isolated using a human CD4+ T Cell Isolation Kit (Miltenyi Biotec #130-096-533) and a manual MACS cell separator (Miltenyi Biotec 130-090-976). Cells were strained using a 40 µm basket cell strainer (Corning #15360801). Cell concentration and viability was determined with a manual hemocytometer counts and Trypan Blue (Thermo Fisher Scientific 15250061) and was 95% or greater for all samples. A 7-cell pool aliquot was thawed, washed, and resuspended in MACS Buffer. 2 million participant-isolated CD4 T cells and 0.04 million 7-cell pool cells were combined, washed, and resuspended in 160 μL D-PBS with 5% FBS (PBS-F) in a Protein Lo-Bind Tube (Eppendorf 0030108116) before proceeding to oligonucleotide-conjugated antibody staining.

### Cell staining using oligonucleotide-conjugated antibodies

10 μL Human TruStain FcX Fc Receptor Blocking Solution (BioLegend 422302) and 4 μL UltraPure Salmon Sperm DNA Solution (ThermoFisher 15632011) were added to cells in PBS-F to prevent nonspecific antibody staining and mixed gently by hand. Sample was incubated on ice for 15 minutes. Lyophilized TotalSeq A human oligonucleotide-conjugated antibody pool (BioLegend 399907) was equilibrated to room temperature and prepared according to the package instructions, with the exception that the antibody panel was rehydrated in 27.5 μL MACS Buffer instead of BioLegend Cell Staining Buffer. 25 μL of the resuspended antibody panel was transferred to the blocked cells, leaving 2.5 μL to prevent precipitates from being transferred. Stained cells were incubated for 30 minutes on ice, with occasional gentle agitation by hand. Cells were washed a total of five times as follows. Cells were washed twice in 5 mL cold MACS Buffer by centrifugation at 500 x g for 4 min at 4°C. Subsequently, cells were washed three times in 3 mL cold MACS Buffer at centrifuged at 500 x g for 4 min at 4°C. On the 5^th^ wash, supernatant was aspirated and the sample was resuspended in 200 µL Cell Resuspension Buffer, consisting of 18% OptiPrep Density Gradient Medium (Sigma-Aldrich D1556) and 0.03% filtered Pluronic F-68 (Gibco 24040-032) in filtered 1x Phosphate Buffered Saline Solution (Gibco 10010-023). Washed cell samples were incubated on ice until ready to load onto the microfluidic encapsulation device

### Microfluidic cell encapsulation and lysis

Cells, lysis buffer, and drop-making oil were prepared in separate syringes. Lysis Buffer consisted of 92.81 mM Tris-HCl pH 8.0 (Corning 46-031-CM), 0.47% Nonidet P-40 Substitute (Roche 11754599001), 5 mM EDTA (VWR E177), and 8 U/μL Proteinase-K (NEB P8107S) in ultrapure water was loaded into a 3 mL Luer-Lok sterile syringe (BD 309657). Cells were resuspended to 3.5 million cells/mL in Cell Resuspension Buffer and loaded into a 3 mL syringe. Dropmaking oil, consisting of 2% 008 Fluorosurfactant (Ran Biotechnologies 008-FluoroSurfactant) in Novec 7500 (“HFE-7500”; 3M Novec 7500), was loaded into a 10 mL Luer-Lok sterile syringe (BD 302995). Syringes were loaded onto syringe pumps (New Era) and connected to the microfluidic device using PE/2 tubing (Scientific Commodities BB31695-PE/2). The cell syringe was kept cooled at 12C. Cells (400 μL/h), Lysis Buffer (400 μL/h), and Dropmaking oil (1600 μL/h) were injected onto the microfluidic device (**Supplementary Fig. 1**) to generate 40μm droplets and the emulsion was collected in PCR tube strips. Dropmaking oil was removed from the bottom of the collection tubes using a syringe or gel-loading pipette tip. Thermocycling oil, consisting of 5% fluorosurfactant (Ran Biotechnologies 008-FluoroSurfactant) in Fluorinert FC-40 (Sigma-Aldrich F9755), was added to the collection tubes so that the total volume of emulsion and oil was less than 100μl. Emulsions were incubated in a thermal cycler for 1 hr at 50°C to perform lysis, followed by 10 min at 80°C to heat-inactivate Proteinase K. After thermal cycling was complete, thermocycling oil was removed using a gel-loading pipette tip or syringe and dropmaking oil was overlaid onto emulsions.

### Microfluidic single-cell barcoding, DNA genotyping, and antibody capture

Cell lysate droplets were merged with PCR reagents and barcoded beads containing poly-T and the forward primer panel. PCR was used to simultaneously attach the bead barcode to the amplicon panel and antibody derived tags (ADTs). The primer panel targeted 10 regions tiled across the HIV genome and 48 additional loci in the human genome. The PCR reagents contained free reverse primers. The reverse primer pool was created by pooling stocks of individual oligonucleotides. 48 human-specific and 10 HIV provirus-specific reverse primers were purchased as individual oligonucleotide stocks from Integrated DNA Technologies (IDT) and pooled to a stock concentration of 100μM HIV primers and 200μM human-specific. A complete list of the forward and reverse primers is found in **Supplementary Table 2**. The PCR solution contained a final concentration of 0.66μM HIV pool and 0.66uM human-specific, 2.76% (v/v) Tween-20 (Sigma-Aldrich), 1.65% (v/v) PEG-6000 (Sigma-Aldrich), 1.65x NEBNext Ultra Q5 Master Mix (NEB M0544S), 0.83 M propane-1,2-diol (Millipore Sigma 398039), 1.24 mM MgCl_2_ (Sigma-Aldrich), 0.25 mg/mL Bovine Serum Albumin (NEB B9000S), in ultrapure water. This was prepared in two steps. First the primers, Tween-20, PEG-6000 and water were combined, heated to 85°C for 5 min, and mixed on an orbital shaker (Eppendorf ThermoMixer F2.0) for 30 min at 37°C. This mixture was combined with the remaining reagents and loaded into the syringe.

The microfluidic merger device consisted of five inlets for cell drops, barcoded beads containing forward primers, PCR master mix with reverse primers, drop spacer oil, and dropmaking oil (**Supplementary Fig. 1**). Syringes for each of these inlets were prepared separately as follows. Five syringes were prepared for the barcoding microfluidic device: two 5 mL syringes (VWR 613-4232) containing drop making oil (one for spacing of cell lysate droplets, the other for generating single-bead droplets), one 1 mL syringe containing Barcoding PCR Master Mix, one 1 mL syringe containing barcoded beads, and one 1 mL syringe containing cell lysate droplets. PE/2 tubing was used to connect all syringes to the barcoding microfluidic device. To run the microfluidic device, cell lysate drops were flowed at 75 μL/h, barcode beads at 150 μL/h, PCR master mix at 275 μL/h, droplet spacer oil at 900 μL/h, bead dropmaking oil at 1100 μL/h. A DC power supply (2V) was connected to a high voltage AC generator (Trek Model 609E-6) connected to an on-chip salt water (1M NaCl) electrode to merge cell lysate droplets and bead droplets. Samples were collected in PCR strips on ice and oil was removed from the collection tubes and replaced with thermal cycling oil. Samples were UV-treated (Analytik Jena UVP XX-15L) for 8 minutes to cleave oligonucleotide barcodes from the beads. Samples were thermal cycled to perform barcoding PCR: 92°C for 6 min (4°C/s ramp); 10 cycles of 92°C for 30 s, 72°C for 10 s, 61°C for 3 min, 72°C for 20 s (1°C/s ramp); 10 cycles of 92°C for 30 s, 72°C for 10 s, 50°C for 20 s, 61°C for 2:40 min, 72°C for 20 s (1°C/s ramp); 72 for 2 min (1°C/s ramp); 4°C hold (4°C/s ramp).

### DNA amplicon and antibody tag recovery

After barcoding PCR, emulsions were broken inside individual PCR tubes by adding 20 μL PFO (1H,1H,2H,2H-Perfluoro-1-octanol, Sigma-Aldrich 370533). A gel loading pipette tip or syringe was used to remove oil from tubes without aspirating the aqueous layer. 50 μL of Tris-EDTA-Tween Buffer (10 mM Tris-HCl pH 7.5, 0.1 mM EDTA, 0.1% (v/v) Tween-20) was added to each tube and tubes were gently agitated by hand to mix, incubated at room temperature for 5 min, and centrifuged on a benchtop microcentrifuge (Eppendorf Centrifuge 5418R). The upper 60 μL of solution containing PCR amplicons was transferred to new PCR tubes, leaving behind sedimented beads. ADTs and DNA amplicons were separated using size selection. Ampure XP beads (Beckman Coulter A63880) were added at a ratio of 0.72:1 beads:sample. The supernatant (containing short antibody tags) was transferred into new PCR tubes. The Ampure XP beads were washed and DNA amplicons eluted in 50 μL nuclease-free water. DNA amplicons were repurified with AMPure XP beads at a ratio of 0.8:1 beads:sample and eluted in 30 μL nuclease-free water. Antibody tags were recovered by adding 1.2 µL of 100 µM biotinylated HDAB 1 (0.6 µM final, **Supplementary Table 2**) and vortexed briefly to mix. Antibody tag tubes were heated to 95 °C for 5 min and snap-cooled on ice. 5μL Streptavidin C1 (Invitrogen 65001) beads per PCR reaction tube were washed in PBS, added to individual tubes to bind antibody tags, washed twice with 200 μL PBS, and resuspended in 30 μL water. All DNA amplicon and antibody tag samples were quantified using a Qubit dsDNA high sensitivity assay kit (Invitrogen Q32851) and Qubit fluorometer.

**Library preparation for next generation sequencing,** <u>DNA amplicons</u>: Libraries of DNA amplicons were prepared using PCR containing 1x NEBNext Ultra II Q5 Master Mix (New England Biolabs M0544S), 0.4 μM standard Nextera i5 and i7 index primers, and 4 ng purified PCR product in a 50 μL total reaction volume. The PCR reaction was thermal cycled to attach P5 and P7 sequences as follows: 98°C for 30 s; 98°C for 10 s, 61°C for 30 s, 65°C for 45 s (12 cycles); 65°C for 2 min; hold at 12°C. DNA amplicon libraries from the same patient run were pooled and purified with two consecutive rounds of 0.7:1 (beads:sample ratio) AMPure XP beads. Libraries were eluted in 100 μL water after the first round and 22 μL after the second round, and quality control (QC) checks were performed using the Qubit and Bioanalyzer 2100 (Agilent, kit #5067-4626). <u>Antibody tags:</u> ADT libraries were prepared using PCR containing 1x NEBNext Ultra Q5 Master Mix and 0.4 µM standard Nextera i5 and TruSeq small RNA N7 index primers (**Supplementary Table 2**), 15 μL of antibody tag samples from the Streptavidin C1 capture reaction in a 50 μL total reaction volume. Antibody samples were thermal cycled per the DNA amplicon PCR recipe, with 15 cycles, pooled, and subsequently purified with two rounds of AMPure XP beads at a bead:sample ratio of 1:1 and 0.85:1 respectively. Antibody libraries were eluted in 100 μL after the first round and 25 μL after the second round, and QC checks were performed with the Qubit and Bioanalyzer. Test sequencing was done to confirm successful library preparation for 8 library tubes on a NextSeq 2000 (Illumina). After successful confirmation, the pooled genomic libraries were combined in batches of five patients (approximately 500,000 cells) at equal amounts and sequenced on a NovaSeq 6000 S4 kit (Illumina) with 150 read 1 cycles, 8 cycles index1, 8 cycles index 2 and 150 cycles read 2. The antibody tag libraries of the corresponding five patients were combined at equal amounts and sequenced on a NovaSeq 6000 S4 kit with 60 read 1 cycles, 8 cycles index1, 8 cycles index 2 and 25 cycles read 2. FASTQ file generation and read demultiplexing were done using the program bcl2fastq (Illumina).

## Bioinformatics pipeline

### Antibodies preprocessing

To convert the paired end fastq files into a unique molecular identifier (UMI)^66, 67^, a corrected cell by antibody count matrix a custom script was used. Paired Fastq files were iterated through entry by entry, and the three cell barcode blocks compared to their barcode white list. For each barcode a maximum of 1 edit distance (Levenshtein distance) ^68^ was tolerated. For entries passing the barcode test, the antibody barcode was compared against its whitelist, tolerating 1 edit distance, to assign the antibody identity. Reads passing both checks were written into a temporary ASCII file in a compressed format analogous to the BUS format^69, 70^. In these files each line is a quadruple of a unique cell barcode identity, antibody identity, UMI code and number of occurrences of the respective combination. To arrive at the final raw cell by antibody matrix, UMIs most likely originating from a PCR error were collapsed according to the “directional” rule using UMI-tools^71^.

### Batch effect removal and cell type annotation

To account for batch effects between experiments, antibody count matrices were further processed with the TotalVI package^23^ (v. 0.17.4) using default settings. TotalVI was designed for CITE-seq experiments^72^ and embeds a transcript count matrix and an associate antibody count matrix jointly in a low dimensional manifold (set to 20 dimensions) through a variational autoencoder. Since DAb-seq experiments yielded antibodies and DNA but no transcript measurements, we provided the method with an all-one dummy RNA vector (to satisfy the input requirements) and ADT data. The RNA component of the model thus contains no information and low dimensional embedding is driven by the antibodies. The low dimensional representation was then used to create a nearest neighbor graph with scanpy (v. 1.9.1)^73^ and UMAP (v 0.5.3)^74^ using default parameters, which was used for unsupervised clustering with the Leiden method^26^ from the python package leidenalg (v. 0.8.10), using a resolution parameter of 3.

To annotate the blood cell types, seed labels were generated by transferring the cell labels from CITE-seq experiments^27, 28^ to our dataset. For this, we restricted the data in both studies to the intersection of antibodies and used scANVI^75, 76^ to transfer the labels. Transferred seed labels were inspected and compared against known antibody markers and, if necessary, manually curated. Finally, all cells within the Leiden clusters were annotated with the most frequent seed label within the cluster to eliminate occasional errors in label transfer.

### Genomic read preprocessing

Cell identification, sequence mapping and variant calling for DNA sequencing reads was done as previously described ^18^ with hg17 as human reference and HXB2 as reference for HIV. Briefly, sequenced reads from the genomic libraries were filtered for correct cell barcodes (analogous to the processing of antibody data), by identifying the 3 barcode blocks and matching them against the whitelist. Reads with no more than one edit distance per block were retained. We mapped the sequences with Bowtie2^77^ against the expected amplicon sequences, which were extracted from both reference genomes. The number of mapped reads for each amplicon and cell forms a cell by amplicon count matrix. We used the marginal cell distribution from this matrix to discriminate cell containing drops from empty drops by the “knee”-method^78^, and call droplets with higher total amplicon count than the inflection point as cell-containing. This is an established heuristic that exploits the fact that cell containing droplets produce an order of magnitude more amplicons than noise in the droplet workflow. To identify spike-in cell lines from their genotype, we used GATK^79^ to call single nucleotide polymorphisms from the aligned human amplicons as described previously^18^. DNA variants were used to differentiate between blood cells from patient samples and the spike-in cell lines.

### Identifying cells with HIV

As J-Lat cells harbor an intact provirus and exhibit a distinct surface immunophenotype compared to PBMC cells, we used the unsupervised clustering of the batch corrected antibody count matrix to separate cells into J-Lat and PBMC and assess false positive and false negative detection in the two subsets. We postulate that two distinct mechanisms can lead to false positives: First, concomitant encapsulation of a HIV infected cell with a non-infected cell (cell doublets) leads to chimeric cell profiles. Such false positives are intrinsic to droplet experiments and controlled through the average cell loading rate (Poisson statistics) which we set at 0.049, yielding an expected doublet and multiplet fraction of 2.4%. Second, ambient counts originate from cell-free DNA and barcode hopping during PCR amplification of the sequencing libraries. In contrast to single cell RNA experiments, premature cell lysis is unlikely to cause cell free DNA as it is compacted into the nucleus. We hence expect barcodes hoping to contribute the majority. To control for barcode hopping we call cells as HIV positive if they have at least two amplicon counts for at least one HIV amplicon.

### Detection of cells with “intact” HIV

To differentiate potentially intact proviruses from defective ones, we assessed the amplicon-specific technical dropout probability using the PBMC, J-Lat spike-in experiment. Our coverage does not include the splice junction at position 743 or the packaging signal at position 680 (HXB2 coordinates), preventing the detection of these specific features. Although this panel cannot detect most small structural defects, large deletions account for over 45% of defects^15^. These large deletions can lead to the loss of target amplicons, which should be detectable as amplicon dropout. Given that the J-Lat provirus is intact, any amplicon that fails to amplify in these cells must be technical. Since technical effects and differences in the viral sequences between experiments may affect dropout probabilities differentially, we do not set an absolute threshold on the likelihood of identifying potential intact viruses. Instead, we classify the top 10% of cells per patient as “intact” and the bottom 33% as “defective” (with cells in between considered ambiguous). Importantly, with a coverage of only 24% of the HIV genome, even in the absence of technical dropout, perfect identification of intact viral sequences is fundamentally impossible. A probabilistic statement is, therefore, the best that can be achieved. We chose a 10% cutoff to balance between the empirical observation that only about 5% of parvoviral sequences are intact^15^ and too many false negatives due to imperfect detection sensitivity for full-length genomes of our assay.

### Differential expression

To investigate differentially expressed epitopes in HIV+ and HIV-cells, or cells with intact versus defective HIV, we restricted the analysis to CD4 T cells. Differential expression analysis of a particular marker between two groups can be formalized as a generalized linear model. Such regression models are intimately related to the T-test and employed in more advanced differential expression models such as DEseq2 or edgeR^80, 81^. Because ADT counts are integers ≥ 0, the basic assumptions of the T-test are not met, and DEseq2 or edgeR, which are dedicated tools for RNA sequencing, employ heuristics that may not be justified for protein data. For this reason, we chose to employ the simplest model appropriate for count data: Poisson linear regression. We model the effect of HIV infection as a two-level hierarchical Bayesian generalized linear model, with infection status at level 1 and participant ID at level 2 to account for technical batch effects and interpersonal variability. Specifically, we implement the following model in NumPyro (version 0.12.1), which we fit for each antibody independently:

Level 1 effect:

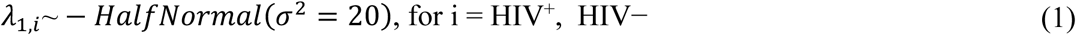

Level 2 Effects:

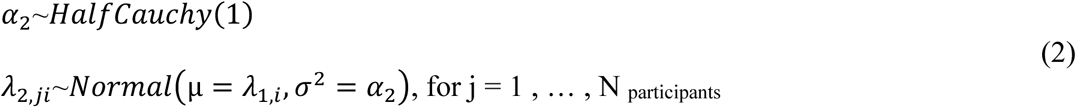

Observation Model:

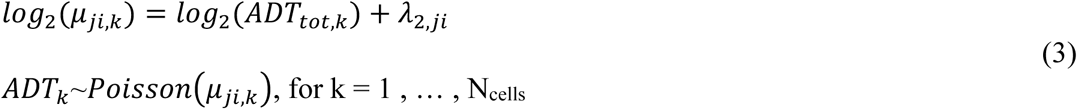

With:

- Level 1 effect *λ*_1,*i*_ modelling antibody expression given HIV status (e.g. 0 = negative, 1 = infected)
- Level 2 effect *λ*_2,*ji*_ modelling batch effects from biological and technical variability given batch index (j = 1, …, 8; note that experimental batch and participant ID are confounded in our experimental setup) and HIV status
- *ADT_tot,k_* represents the sum of all observed ADT counts for cell *k*. Note that for each cell *k* the indexes *j,i* are known, hence we dropped them from the final relation.

For each antibody, we estimate the model parameters using Markov Monte Carlo (MCMC) sampling with the NumPyro implementation of the No U-turn Sampler (NUTS)^82^. From the MCMC samples, we obtain the posterior distribution for the parameter λ_1_, which represents the effect size for HIV infection on the expression profile of the ADT. To represent the data as volcano plots we use the mean of the posterior distribution as effect size and the type S-error to represent statistical significance^83^. We compute the type S-error for positive effects as:

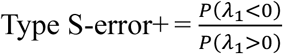

We compute the type S-error for negative effects as:

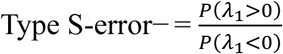

The overall type S-error is the maximum of the two values:

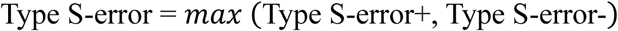

Where:

*P*(*λ*_1_ > 0) be the proportion of posterior samples where λ_1_ > 0.

*P*(*λ*_1_ < 0) be the proportion of posterior samples where *λ*_1_ < 0.

All code for the differential expression calculation can be found in the paper associated GitHub repository (**Data Availability**).

### Accounting for cell type effects

While it would be straightforward to extend this model to include effects caused by different cell types via a third indicator variable (which could be crossed or hierarchical), we chose to correct it via data stratification. The rationale is that data stratification does not require an explicit linear model approach and can be applied to simpler differential expression approaches such as T-tests or the procedure proposed by Sun et al. ^19^. The goal of the procedure is to remove the correlation between HIV infection and distinct cell subsets. To do this, we retain all observed HIV-infected cells and identify the subset with the highest rate of infection. We then iterate through each cell subset and randomly select and discard non-infected cells until the rate of infection within the subset reaches the same frequency as in the most highly infected subset. The resulting cell sample is the stratified data set.

### Reproduction of differential expression analysis in Sun et. al

We replicate the differential expression approach in Sun et al.^19^ by performing the following steps: We convert the ADT count matrix to central log ratio (CLR) transformed data for each cell. The CLR uses the geometric mean in the denominator, and it is undefined for zero counts, requiring the addition of a pseudo count. Next, we convert the CLR transformed data to binary (not expressed, expressed) by setting the matrix elements to 0 if they are smaller than the empirical threshold of -0.104 (personal communication), and to 1 otherwise. To account for differences in the number of observed HIV+ (or intact) cells between participants, we replicate the authors’ proposed bootstrap procedure. This procedure samples with replacement an equal number of HIV cells from each participant such that the total number of sampled HIV cells equals the sum of all observed HIV cells across patients. This means that some HIV cells are counted multiple times, guaranteed in participants with fewer than average infected (or intact) cells, and not all cells are used, particularly in participants with more than average numbers. Similarly, for the comparator group, we sample an equal number of non-infected (or defective) cells from each patient, ensuring the total sampled cell number matches the number of measured cells. The goal is to weight each participant equally, but this procedure discards cells from participants with high infection frequency while counting cells from participants with lower infection frequency multiple times, thereby systematically decreasing the variance of the HIV expression profile. We calculate fold enrichment directly from the resulting contingency table describing the number of occurrence of marker positive and negative cells in the HIV positive and negative groups, and perform a χ2-test to assess statistical significance. To investigate the stability of this procedure, we repeat it 250 times and report the mean value of the observed fold change and p-value distribution and denote one standard deviation by the error bars.

### Odds ratio calculation

To test for variability in the cell enrichment we use binomial regression.

We model the probability of cell being infected, given participant id, and cell type as:

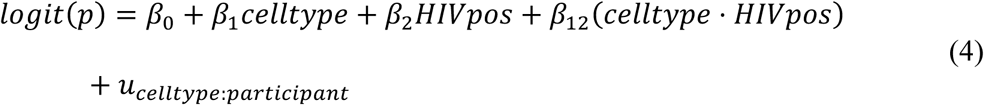

Where:

- *β*_0_ is a global offset
- *celltype* is an indicator vector denoting the cell subset membership for each cell and *β*_0_ are the associated coefficients.
- *HIV_pos_* is a binary vector denoting cells that are infected (or intact)*β*_2_ are the associated coefficients.
- *β*_12_ are the coefficients for the fixed effect interactions between cell type and HIV infection
- *u_celltype:participants_* ∼*Normal*(0,σ^2^) is a random effect term modeling the interactions between participants and cell types.

From the fitted model we calculate the odds ratio of being infected:

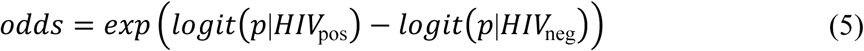

We implement this linear regression model as Bayesian model in python bambi (v 0.13.0) using the default priors. We report the mean, 5 percent percentile and 1 percent percentile of the posterior odds ratio distribution. We proceed analogously for the stratified data set. The python implementation is available from the paper associated GitHub repository (Data Availability). Note that the log odds calculation involves normalizing against the total number of infected cells of a particular class (LTR pos, 0-30 percentile, 30-60 percentile etc.) and hence the directionality of change is lost. For example, a positive change in HIV intact cells within the Naive subset could also result from a decrease in the proportion of HIV intact cells in other subsets.

## Contributions

CLD, ICC, and ARA conceptualized experiments. SGD, EAB, and MS recruited and organized sample collection for the study. CLD, SS, KMJ, YP, BD, and ICC performed experiments. CLD, ARA, and ICC wrote the manuscript.

## Acknowledgments

ARA and CLD were supported by CZ Biohub. ICC was supported by grants from the NIH (K22AI152644, 1R01DA059551-01). EB was supported by the Intramural Research Program of NIAID (ZIA AI005131). A Collaboration for AIDS Vaccine Discovery (CAVD) grant from the Bill & Melinda Gates Foundation (INV-008500) supported the Reservoir Assay Validation and Evaluation Network (RAVEN) Study.

---

**Supplemental Figure 1.**
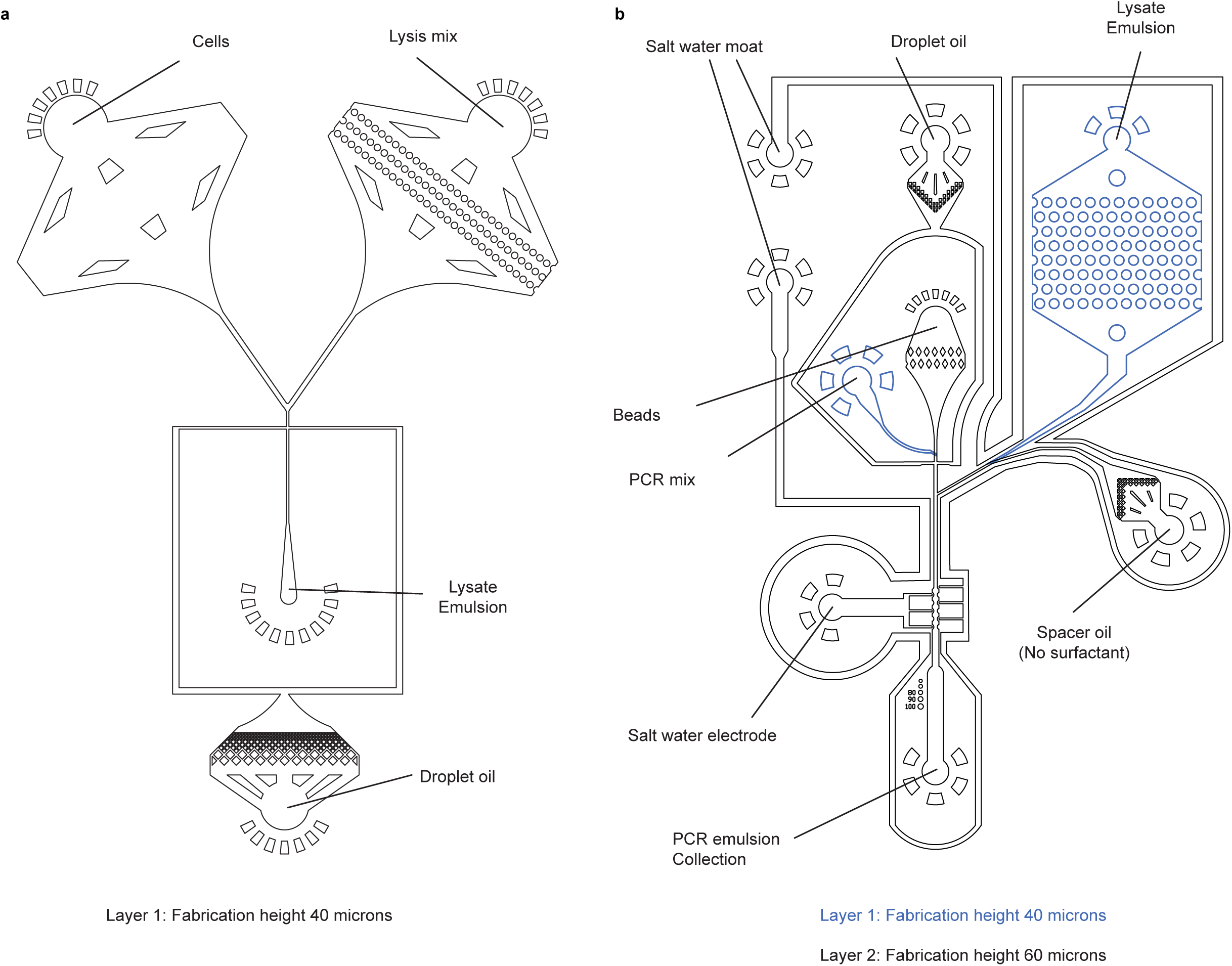
Schematics of the Two Microfluidic Devices Used in this Study. **a)** Coflow device designed for cell encapsulation with inlets and outlets indicated. **b)** Optimized two-layer merger device designed for this study with inlets and outlets indicated.

**Supplemental Figure 2.**
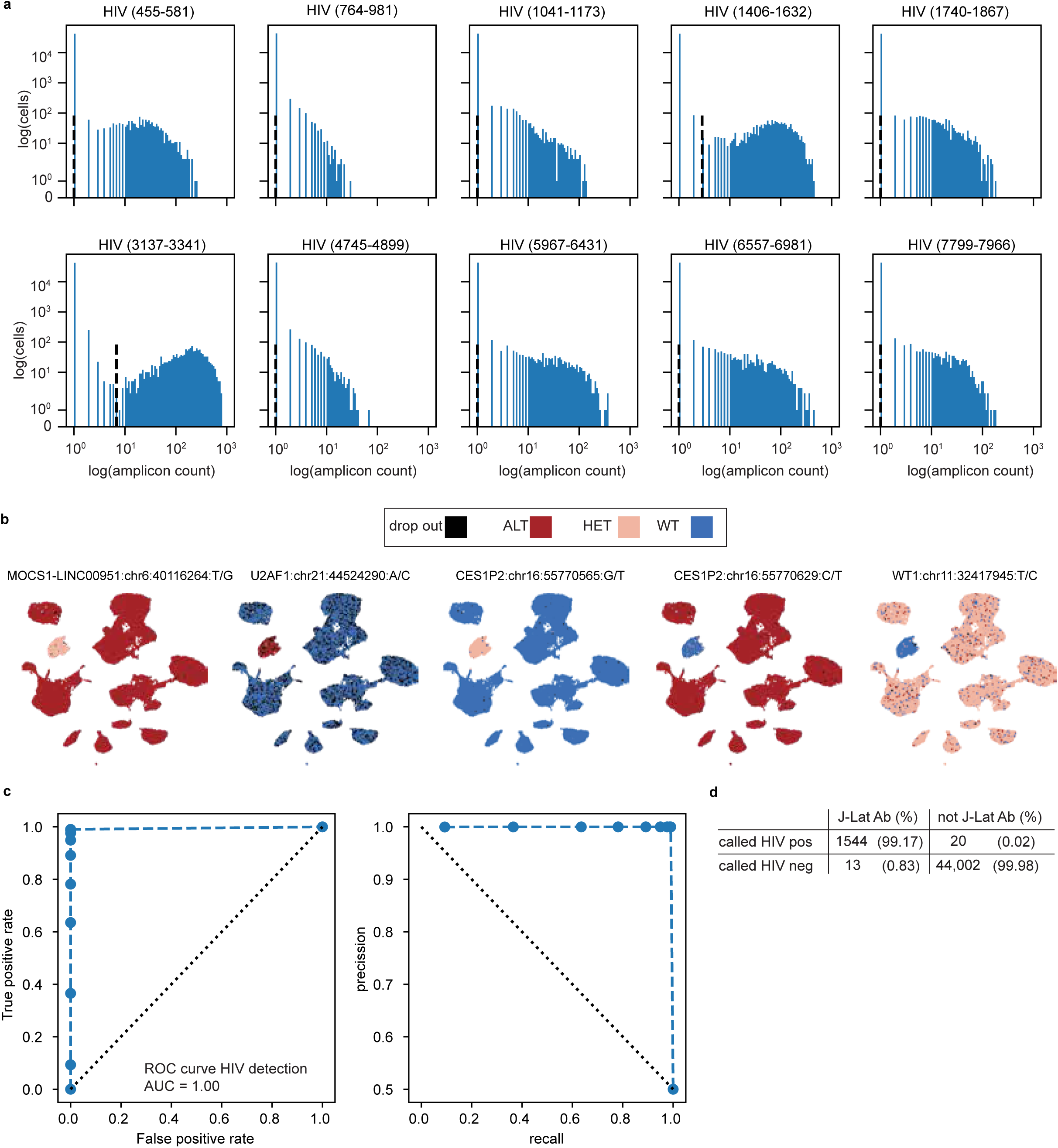
Performance of Single-cell HIV DNA Detection in J-Lat PBMC spike-in data. **a)** Amplicon count distribution for the 10 HIV amplicons and J-Lat cells (see also summary plot in Fig. 3d). Target specific differences in the amplification efficiency are apparent. **b)** Single nucleotide polymorphisms (SNPs) that distinguish J-Lat cells from the HIV-PBMC donor overlayed on UMAP plots. Five genomic variants were used to differentiate between the two donors. Black indicates target dropout; blue, wild type heterozygous; rose, heterozygous, and red homozygous alternate. **c)** Receiver operator characteristic (ROC) and precision recall curves for HIV genotype detection in the J-Lat cluster. **d)** Confusion matrix between genomic HIV detection (rows) and J-Lat phenotype (columns). Absolute cell counts and percentages are given in parentheses.

**Supplemental Figure 3.**
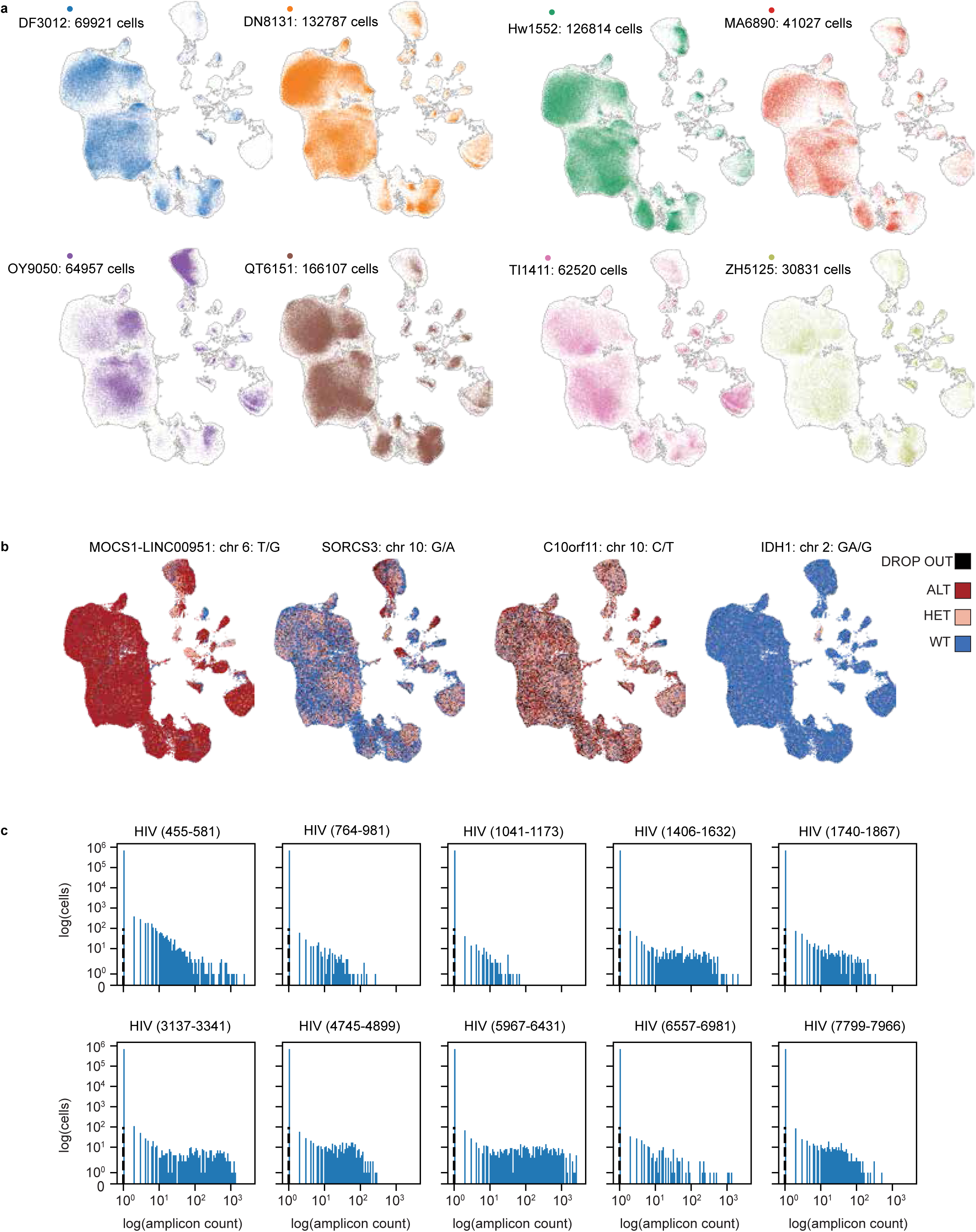
Cell and Amplicon Distribution in Samples from PLWH. **a)** Integrated UMAP across all sequenced cells from all participants is shown as a light outline. Independent participant samples are shown in color to illustrate the sample coverage (see Supplemental Figure 4 for combined data). **b)** A selection of four genomic variants that differentiate between study participants and spike in cell lines. The precise genomic coordinates are withheld to protect the identity of the study participants. **c)** HIV amplicon count distributions from all samples from PLWH.

**Supplemental Figure 4.**
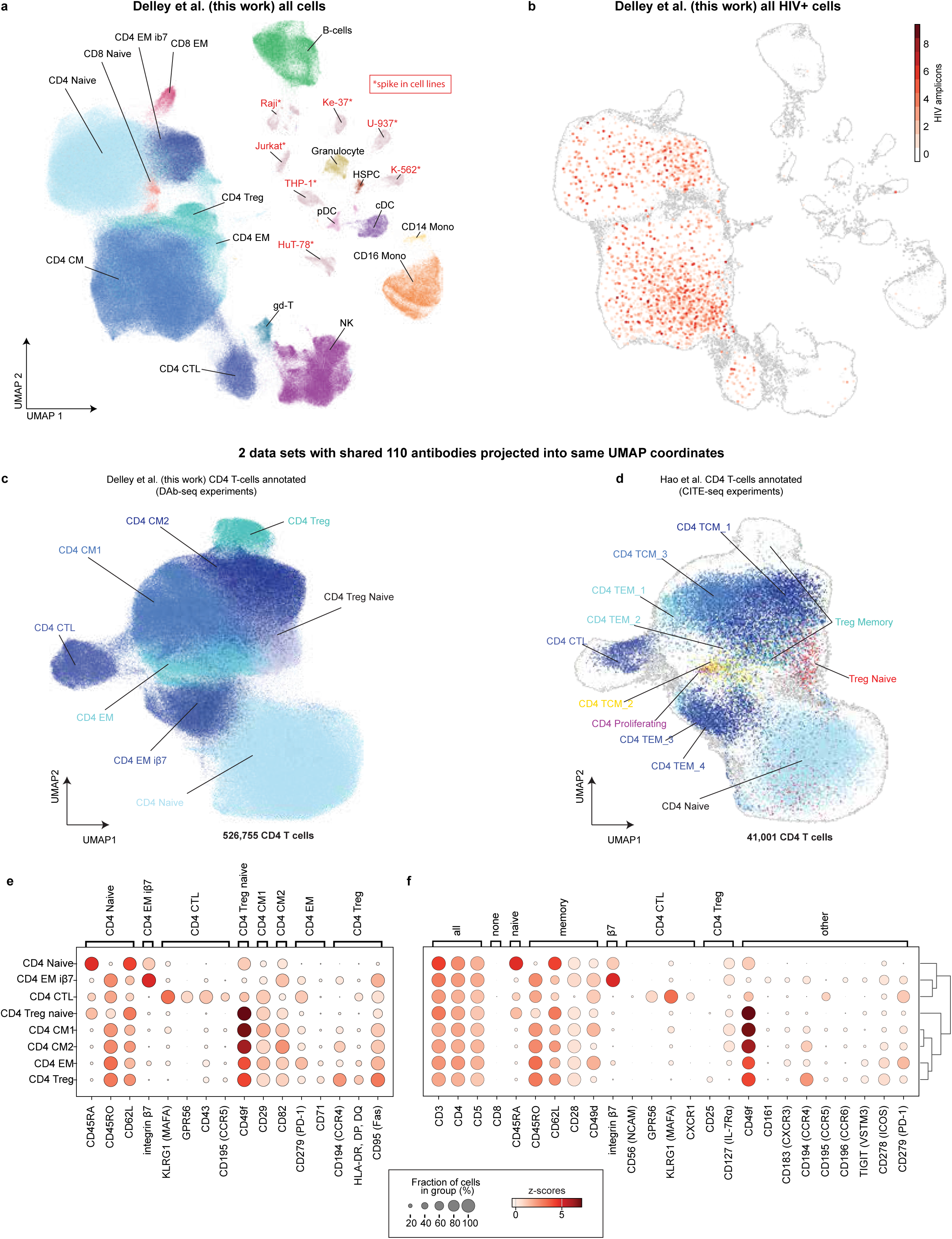
Antibody Markers Delineating Cell Types. **a)** UMAP of DAb-seq data from this study, annotated with cell type labels. Clusters from spike-in cell lines have red labels. **b)** Location of HIV+ cells within clusters. **c)** Integrated UMAP of DAb-seq data annotated with label transfer from CITE-seq^28^ using 110 shared antibodies. **d)** Integrated UMAP with data from Hao et al. **e-f)** Markers used in manual annotation of cell types. Dot plots depict marker expression in CD4 T-cell subsets. Dot size indicates the number of cells expressing the marker, and shading represents the mean Z-scores (normalized across cells).

**Supplemental Figure 5.**
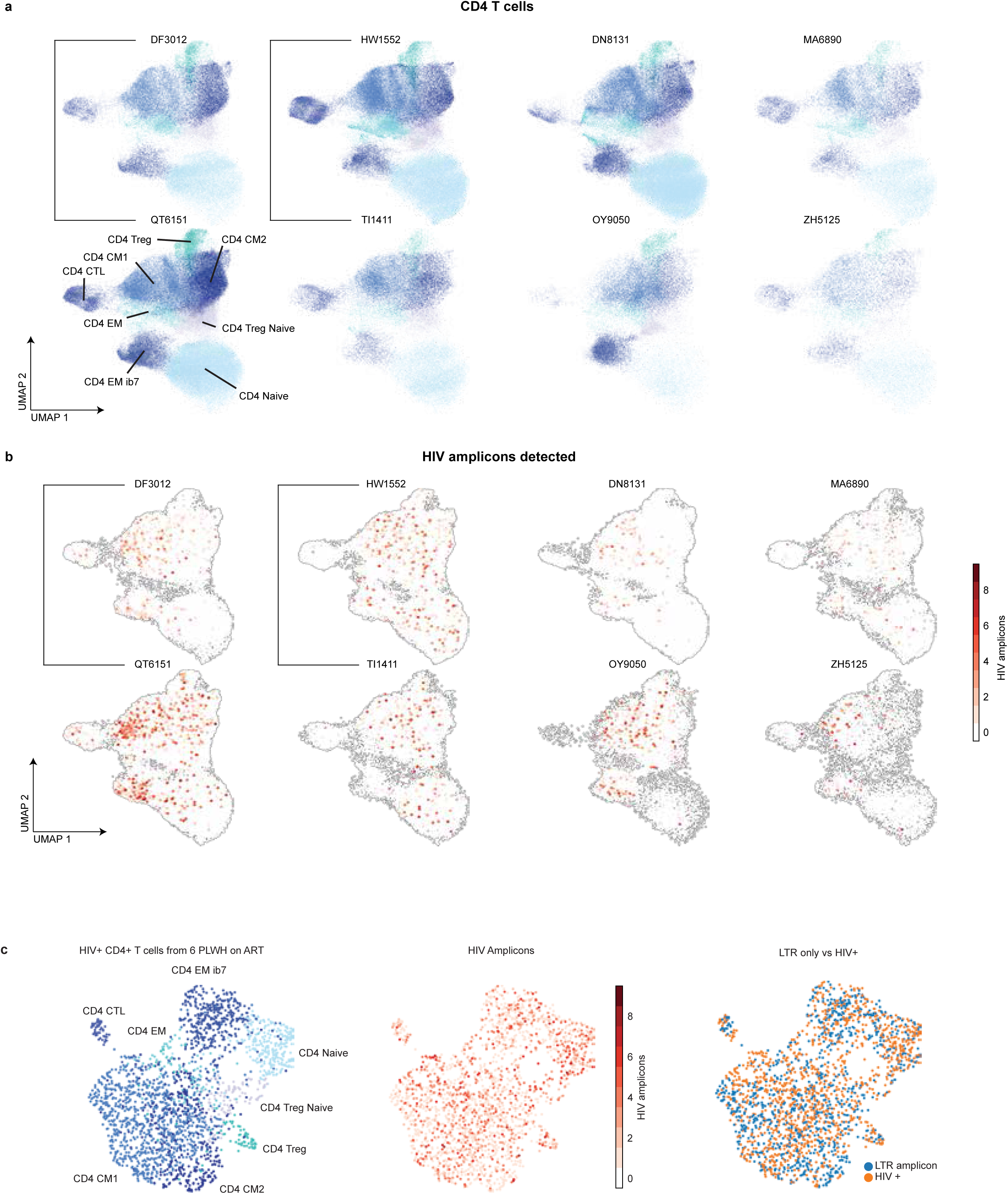
UMAPs of All Detected Cells from PLWH Resolved by Participants. **a)** UMAPs of data from individual samples. The integrated UMAP of all CD4+ T-cells is shown as a light outline. **b)** Location of HIV+ cells within clusters. The sample pairs DF3012, QT6151, and HW1552, TI1411 are different time points from the same person **c)** UMAP of only HIV+ cells colored by cell subtype (left), the detected number of HIV amplicons (middle), and LTR-only amplicons (right).

**Supplemental Figure 6.**
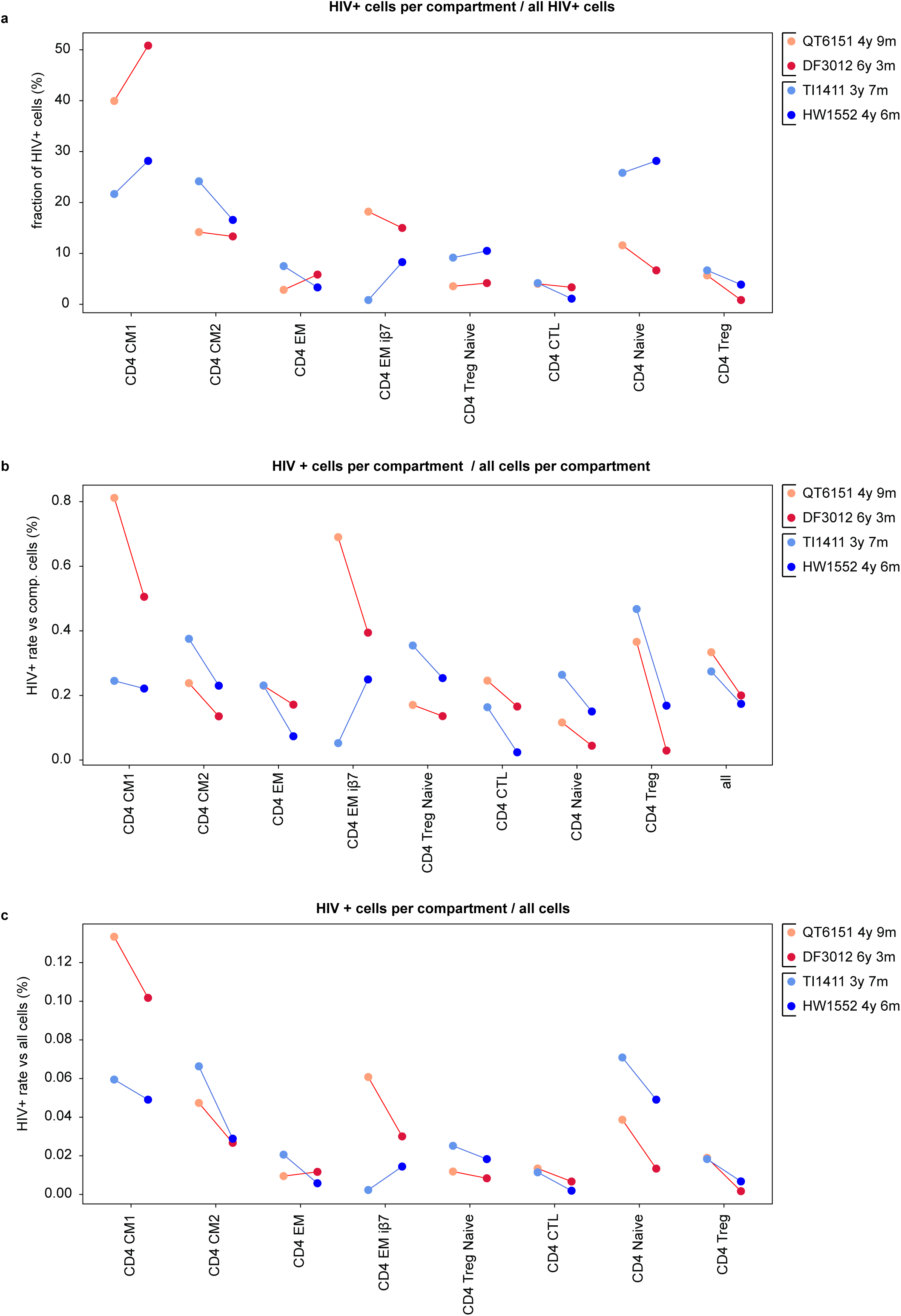
Analysis of Longitudinal Samples. a-c) Change in the abundance of HIV+ cells within CD4 subsets for two participants at two time points. **a)** Fraction of HIV+ cells in cell subsets per all HIV+ cells. **b)** Fraction of HIV + cells in subsets per all cells in a subset. **c)** Fraction of HIV + cells in subsets per all cells.

**Supplemental Figure 7.**
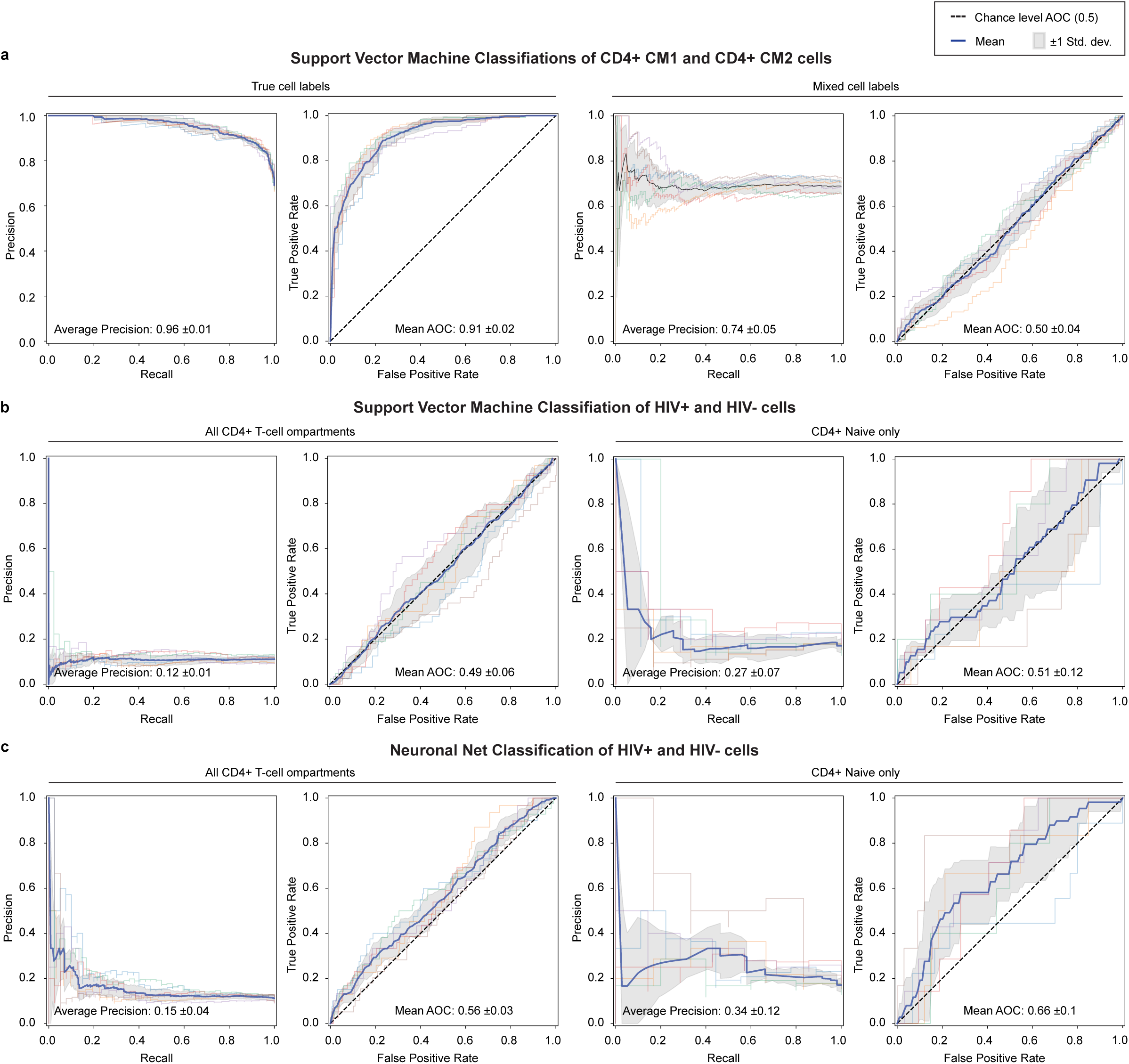
Linear And Nonlinear Classifiers to Identify Marker Combinations that Predict Infection Status. **a)** Support Vector Machine Classifications of CD4+ CM1 and CD4+ CM2 cells. Left two panels give precision-recall (PR) and receiver operator curve (ROC) for a linear Support Vector Machine (linear classifier) trained to classify CM1 and CM2 data. The training data was split for 6-fold cross-validation (faint lines). The solid line indicates the mean classification across cross-validation, and the shaded gray area is one standard deviation. The dashed line indicates chance level predictions. Right two panels, the same classifier trained on the same data set with randomly permuted cell type labels. As the permuted labels are not informative, as expected, the prediction is not better than chance. **b)** Support Vector Machine Classification of HIV+ and HIV-cells. Left two panels, the same classifier and analogous display as in a) but trained on HIV+ versus HIV-cells across all CD4 T-cell subsets. Right two panels, same as left but trained only on the HIV+ and HIV-cells from the CD4+ Naive subset. Plot are representative of other CD4+ T-cell subsets. **c)** Neuronal Net Classification of HIV+ and HIV-cells. Left two panels, HIV+ vs HIV-across all CD4+ subsets. Right two panels, CD4+ Naive only.

**Supplemental Figure 8.**
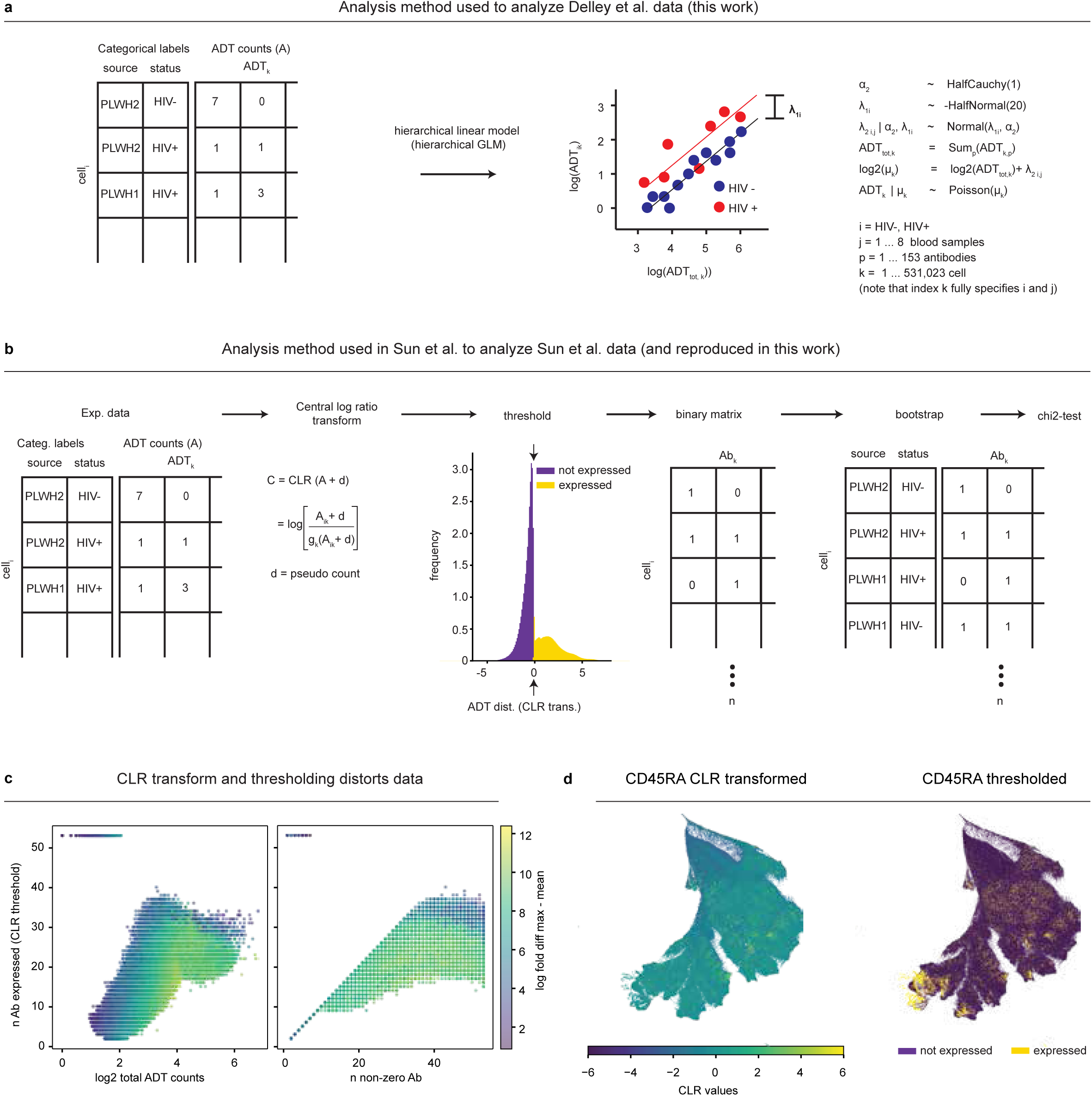
Comparison of the Bayesian Differential Expression Method Used in This Study with the Method Used in Sun et al. **a)** Cartoon representation of the Bayesian hierarchical generalized linear regression used to perform differential expression analysis in this study (see methods). **b)** Cartoon representation of the central log ratio transformation (CLR), binary thresholding, and Chi-squared test used in Sun et al. **c)** The effect of CLR transformation and thresholding on the relationship between the number of antibodies expressed and antibody-derived tag (ADT) counts. **d)** Example of the effect of CLR transformation and thresholding on CD45RA. CLR distorts which cells are “CD45RA+” and shows non-correspondence between count data and binary expression labels.

**Supplemental Figure 9.**
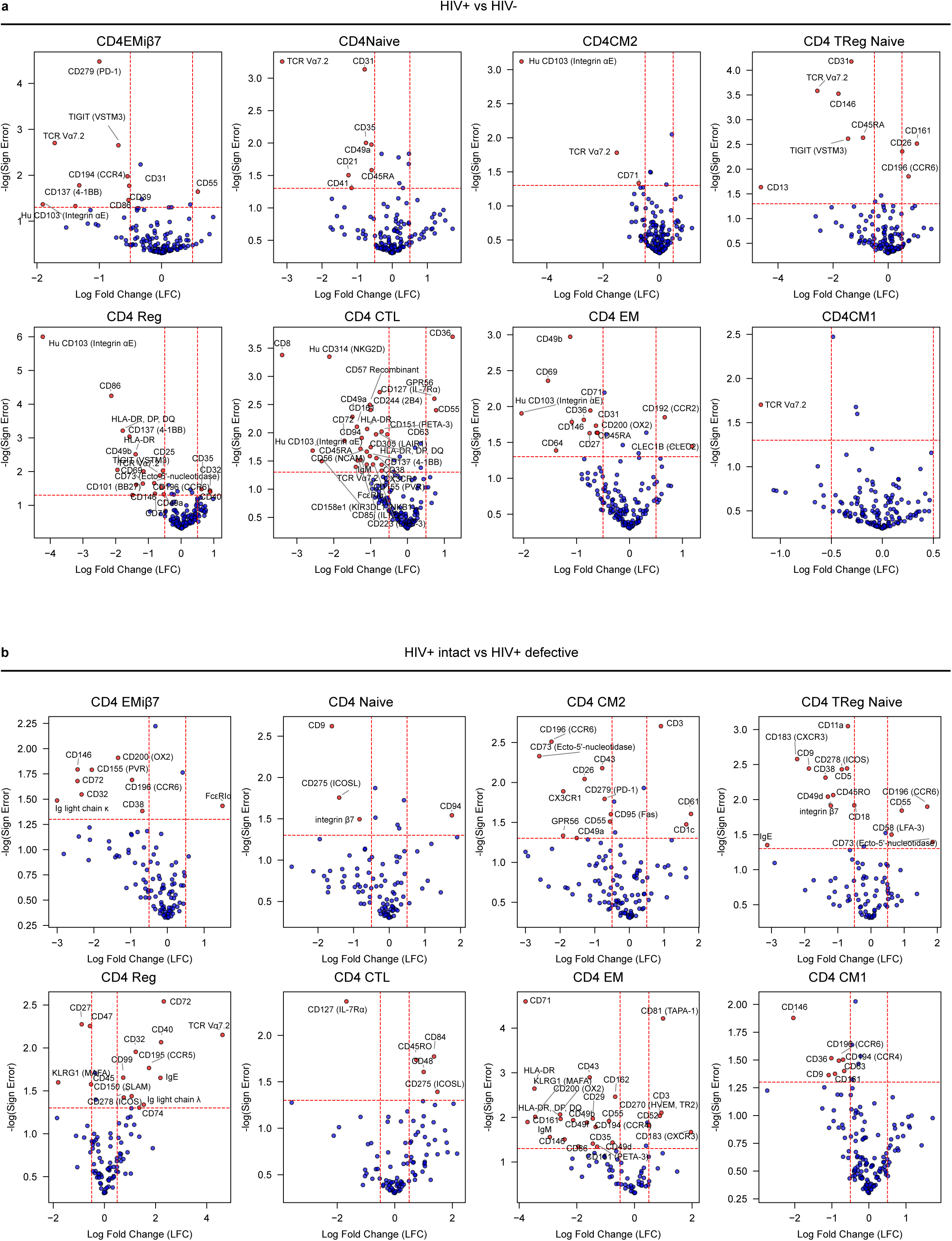
Within CD4 subset Differential Protein Expression (DE) Analysis. **a)** DE between HIV+ and HIV-cells for each CD4 T cell subtype. **b)** DE between HIV+ “intact” and HIV+ defective for each CD4 T cell subtype.

**Supplemental Figure 10.**
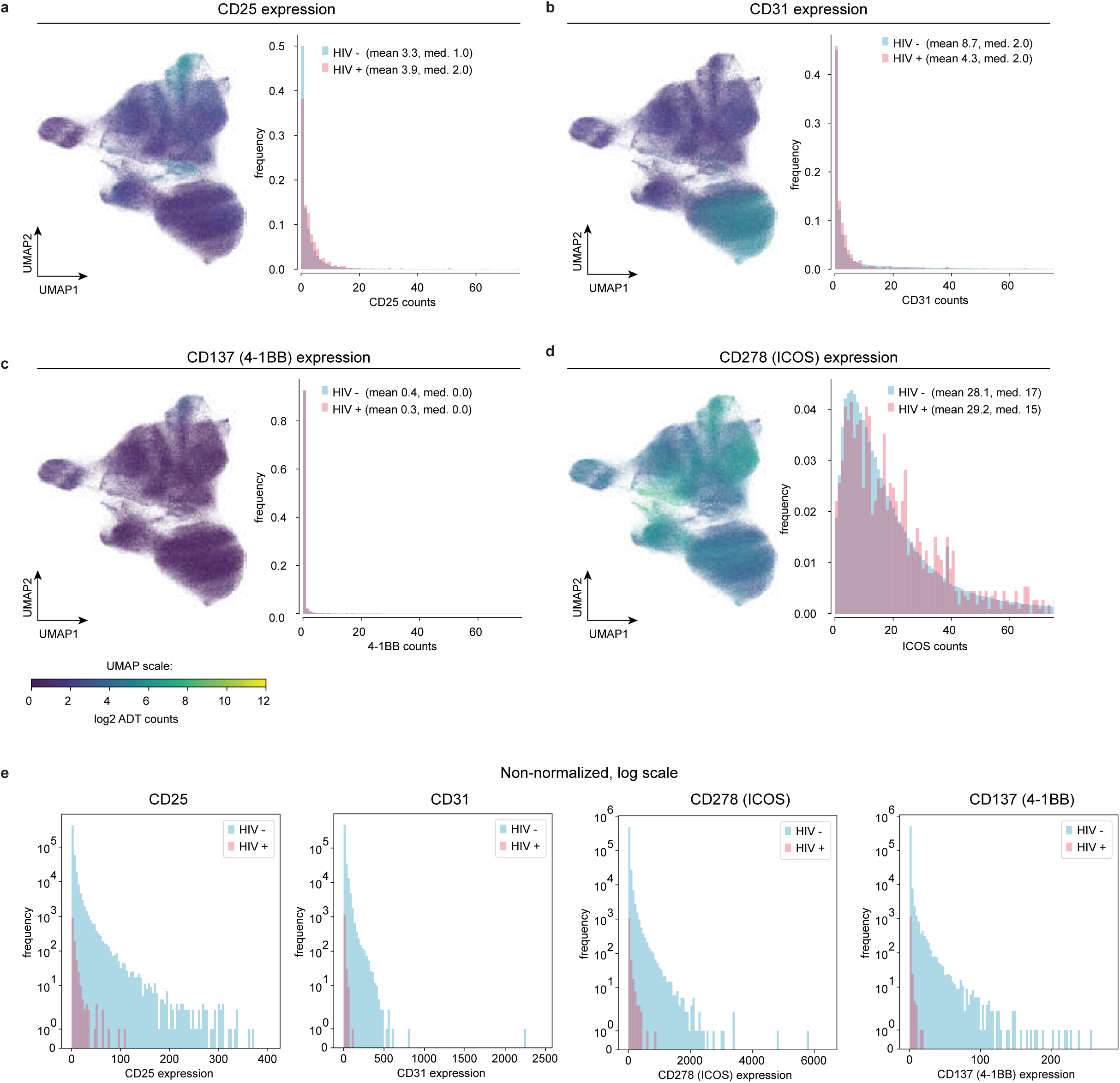
Antibody Expression for Significant (<5% sign error threshold) Proteins after sample stratification. **a-d)** Left, UMAP plots of all CD4+ T-cells with coloring according to Ab signal (log_2_ scale). Right, Ab count histograms for HIV+ and HIV-cells, each group normalized to an area of one. The x-axis is truncated at 75 counts. **e)** Histograms of non-normalized protein expression (log-scale) colored by infection status.

**Supplemental Figure 11.**
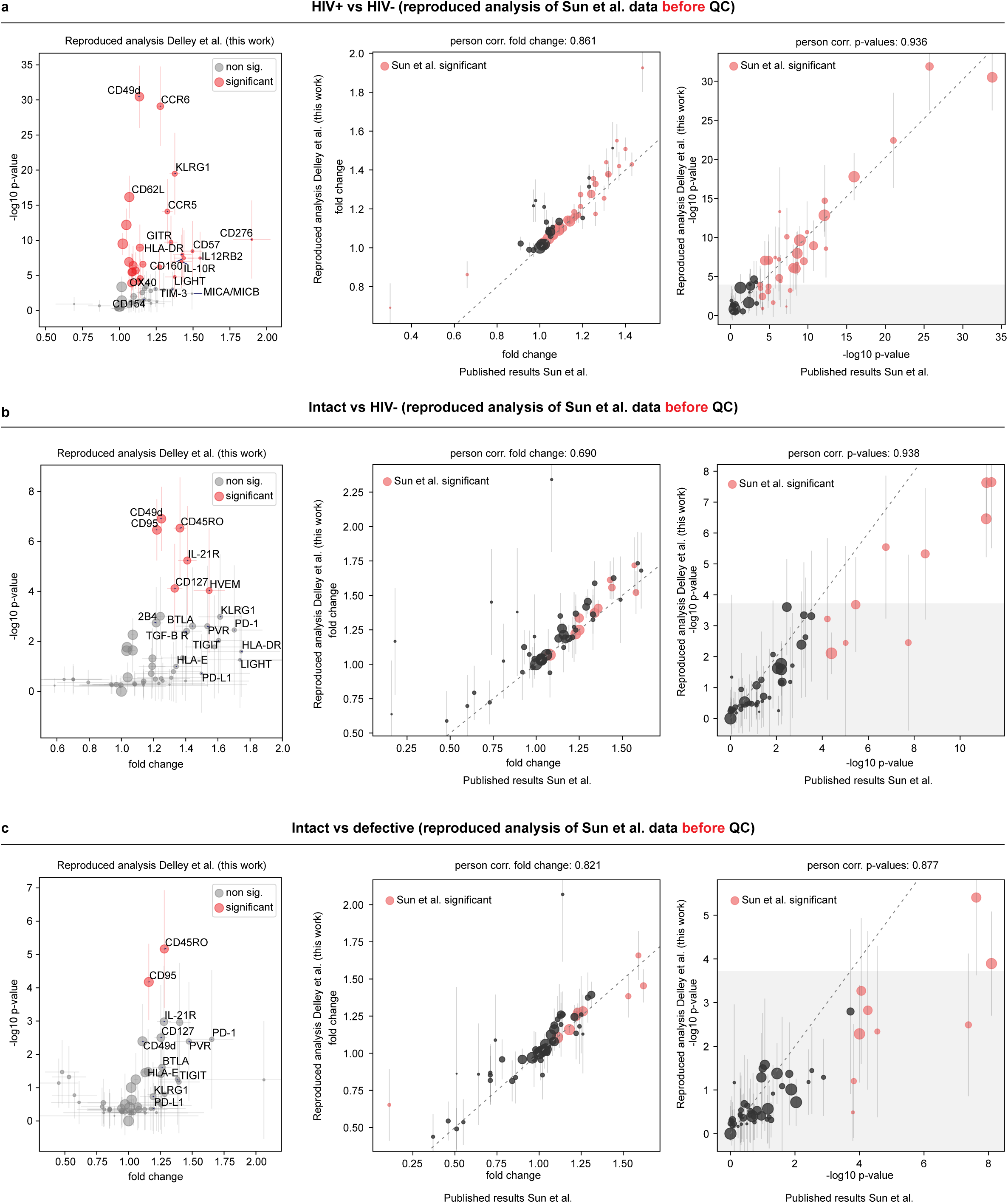
Reproduction of the Differential Expression Analysis from Sun et al. Before Quality Control. **a-c)** Left, Volcano plot of the reproduced DE analyses. Colored dots are colored red if significant at p<0.05 with Bonferroni correction. Middle, Correlation between reproduced (this study) and published^19^ fold changes. Right, Correlation between reproduced (this study) and published^19^ p-values. Middle and Right, colored dots were reported as significant in Sun et al. ^19^ **a)** Differential expression between HIV+ and HIV-cells. **b)** Differential expression between HIV+ intact and HIV-cells. **c)** Differential expression between HIV+ intact and HIV+ defective cells.

**Supplemental Figure 12.**
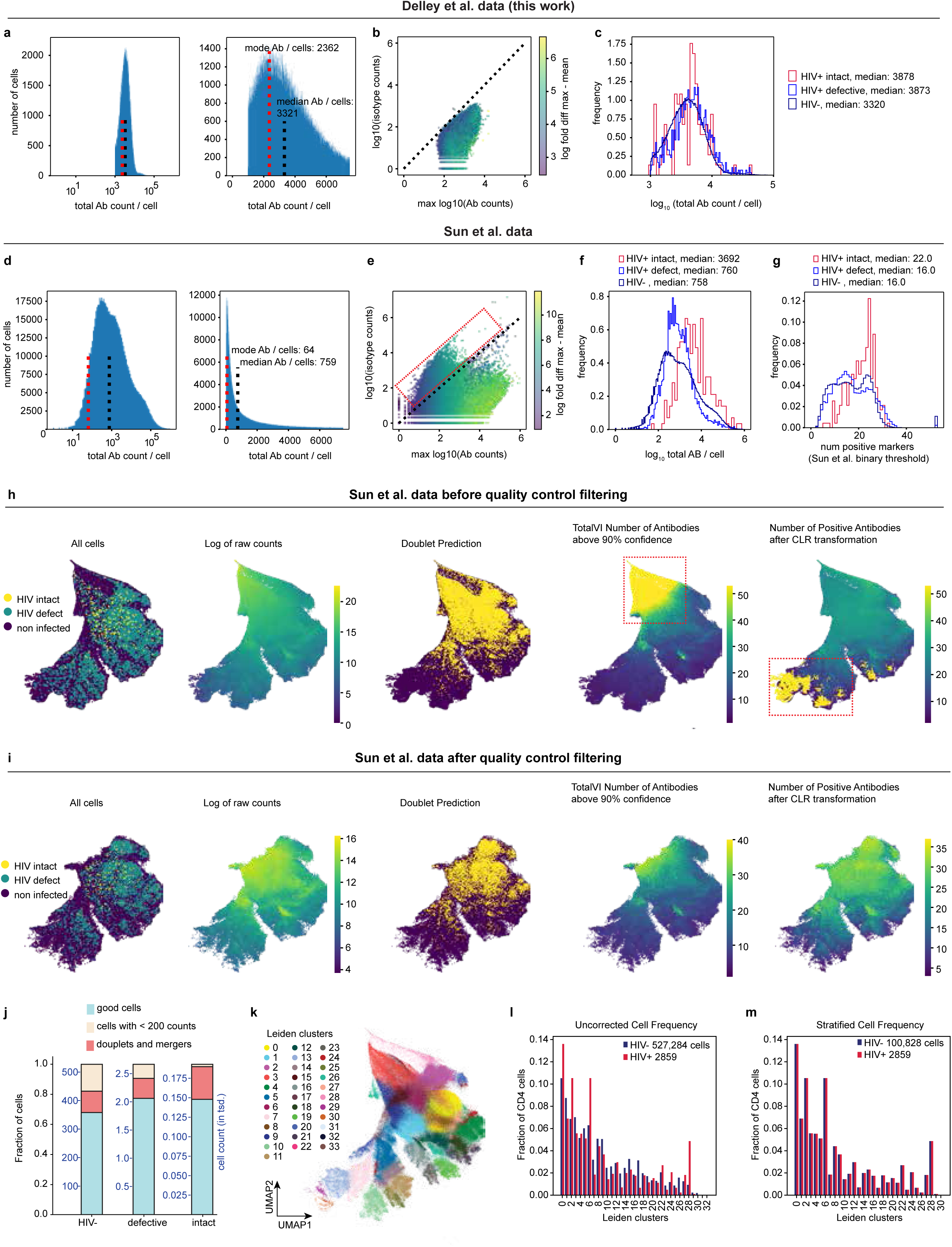
Data Quality Plots. **a-c)** Quality control plots for this study. **a)** Histogram of the total ADT counts per cell. Cells with less than 1,000 UMIs were discarded. Right panel is a zoomed onto the range 0 - 7500. **b)** Total isotype control counts per cell versus the maximum ADT counts per cell. The lower triangle is a conservative estimate of cells with signal-to-noise > 1. The estimate is conservative because cells in the lower triangle have at least one ADT that exceeds the sum of all isotypes and are thus also guaranteed to exceed the count in the corresponding isotype controls. Color scale denotes log_2_ maximum ADT count per cell minus mean ADT count per cell. **c)** Histogram of ADT expression, colored by cells with intact, defective, or no provirus. **d-f)** Quality control plots for Sun et al.^19^ data. **d)** Histogram of the total ADT counts per cell. Data is dominated by droplets with less than 100 ADT counts per cell while at the same time featuring rare droplets with >100,000 counts. **e)** Total isotype control counts per cell versus the maximum ADT counts per cell. High isotype control counts are present in a large fraction of cells. **f)** Histogram of ADT expression, colored by cells with intact, defective, or no provirus. Cells with intact genomes have significantly higher total antibody counts, suggesting technical artifacts. **g)** Histogram of the number of positive ADT makers after Sun et al. binary CLT a binary threshold. **h)** UMAP for Sun et al. data, colored (from left to right) by infection status, total log_2_ ADT count, doublet prediction, number of expressed markers (TotalVI method), and number of positive markers (Sun et al. thresholding). **i)** same plots as h) after quality control steps that remove poor quality cells and doublets. These steps were not performed in Sun et al.^19^. **j)** Bar plot showing the proportion of cells classified as good, low ADT counts, or doublets, as a function of HIV-, HIV+ defective, and HIV+ intact. Cells removed due to low ADT counts (beige), due to indication of mergers or doublets (coral), and retained cells (light blue) are indicated. **k)** Unsupervised Clustering and Data Stratification on Sun et al. data colored by unsupervised clusters (Leiden^26^ method). Cell clustering and labels were not provided in Sun et al. **l, m)** Bar charts depicting frequency of HIV infected and non-infected cells in the respective Leiden clusters before (l) and after (m) after our data stratification to control for the cell types.

**Supplemental Figure 13.**
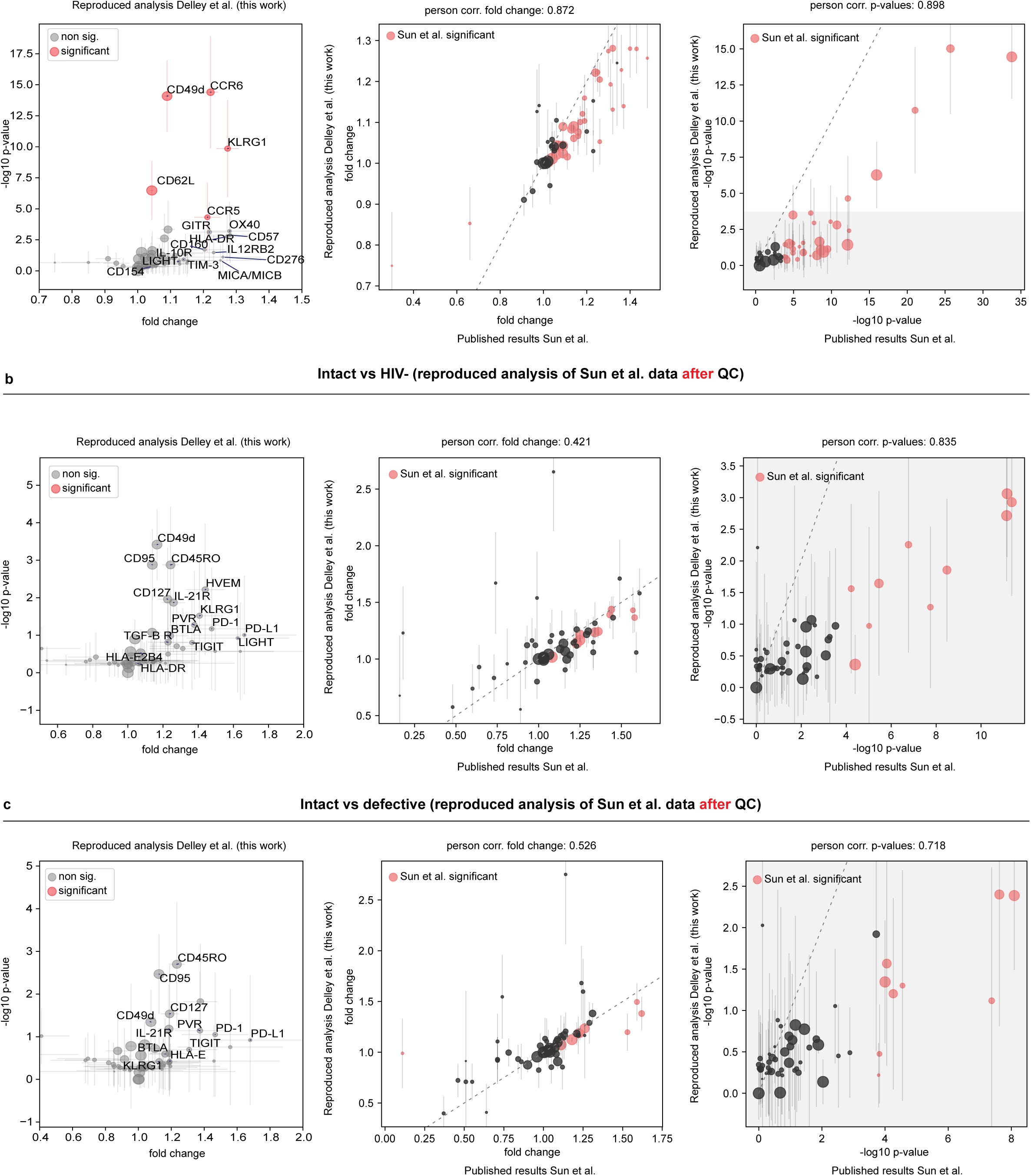
Replication of the Differential Expression Analysis from Sun et al. After Quality Control. **a-c)** Left, Volcano plot of the reproduced analyses. Colored dots in are red if significant at the p<0.05 with applied Bonferroni correction. Middle, Correlation between reproduced (this study) and published^19^ fold changes. Right, Correlation between reproduced (this study) and published^19^ p-values. Middle and Right, colored dots were reported as significant in Sun et al.^19^ **a)** Differential expression between HIV+ and HIV-cells. **b)** Differential expression between HIV+ intact and HIV-cells. **c)** Differential expression between HIV+ intact and HIV+ defective cells.

**Supplemental Figure 14.**
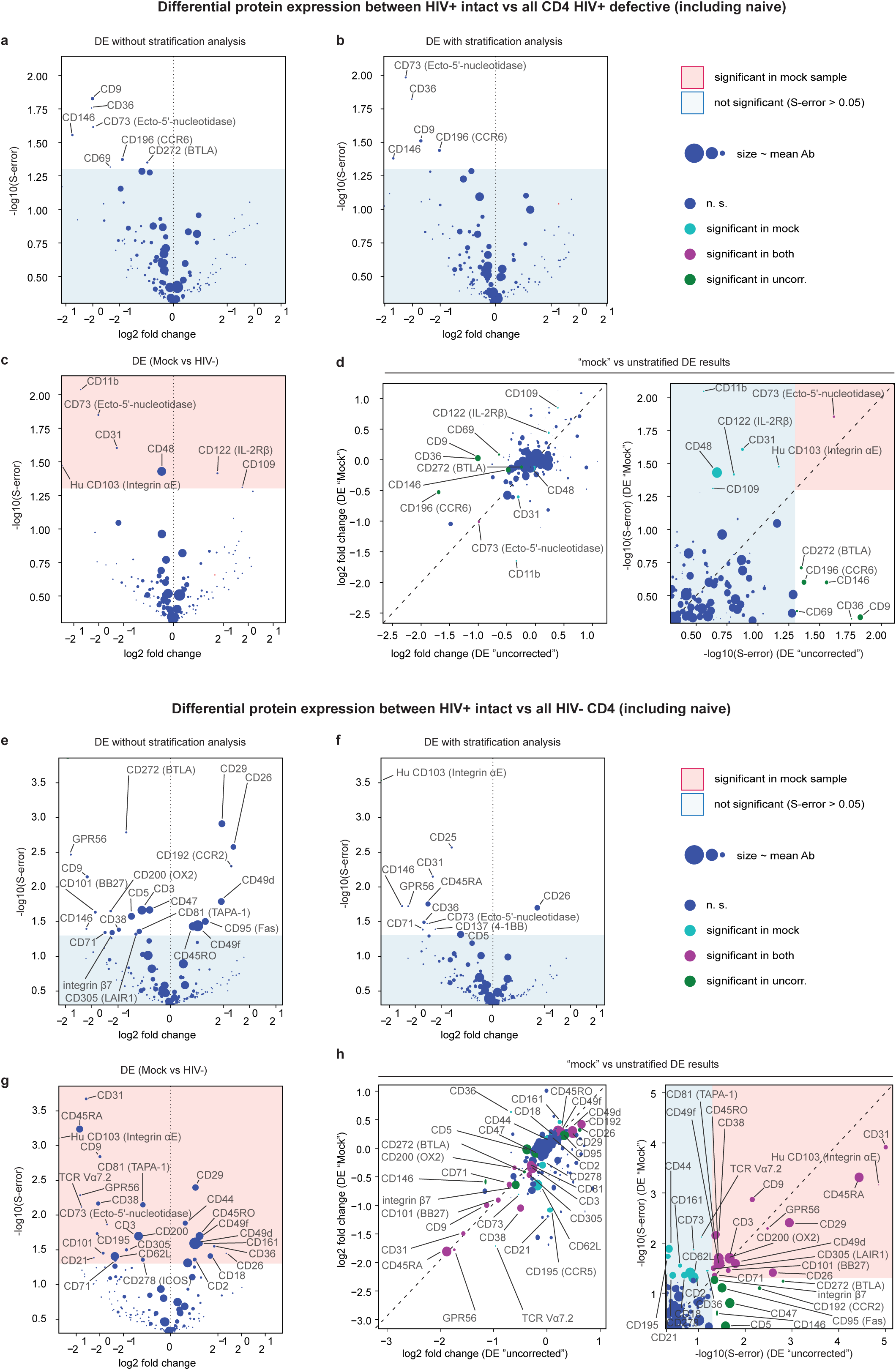
Differential Expression (DE) Analysis for CD4 Cells with “Intact” Versus “Defective,” and “Intact” vs HIV- (This Study). **a-c)** Volcano plots showing DE analysis between CD4 cells with **HIV+ intact vs HIV+ defective** (including naive). **a)** Without cell stratification. **b)** With cell stratification **c)** With mock comparator.) **d)** Correlation of Log fold change (left) and s-error (right) between DE results for the unstratified and mock analyses. **e-g)** Volcano plots showing DE analysis between **HIV+ intact vs HIV-** cells (including naive). **e)** Without cell stratification. **f)** With cell stratification. **g)** With mock comparator. **h)** Correlation of Log fold change (left) and s-error (right) between DE results for the unstratified and mock analyses. **a-h)** “S-error” is the posterior probability of assigning the wrong sign to the effect and is equivalent in interpretation to a p-value.

**Supplemental Figure 15.**
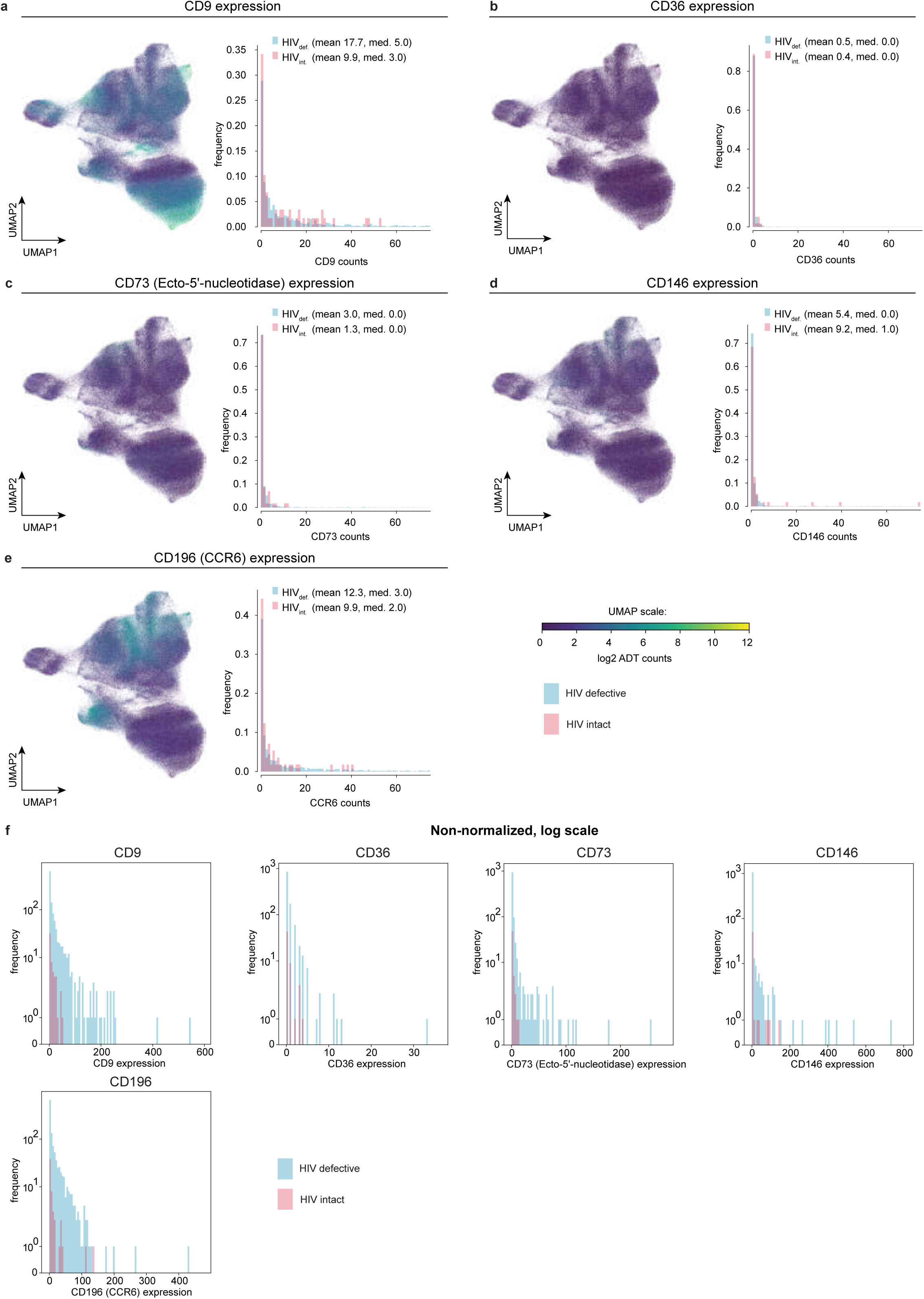
Antibody Expression for Significant (<5% sign error threshold) Proteins after HIV+ “intact” vs. HIV+ “defective” (sample stratification DE approach, this study). **a-f)** HIV “intact” versus “defective” provirus comparison (see Supplemental Fig. 14b). **a-e)** Left, UMAP plots of all CD4+ T-cells with coloring according to Ab signal (log_2_ scale). Right, Ab count histograms, each group normalized to area of one. X-axis is truncated at 75 counts. **f)** Histograms of non-normalized protein expression (log-scale) colored by infection status.

**Supplemental Figure 16.**
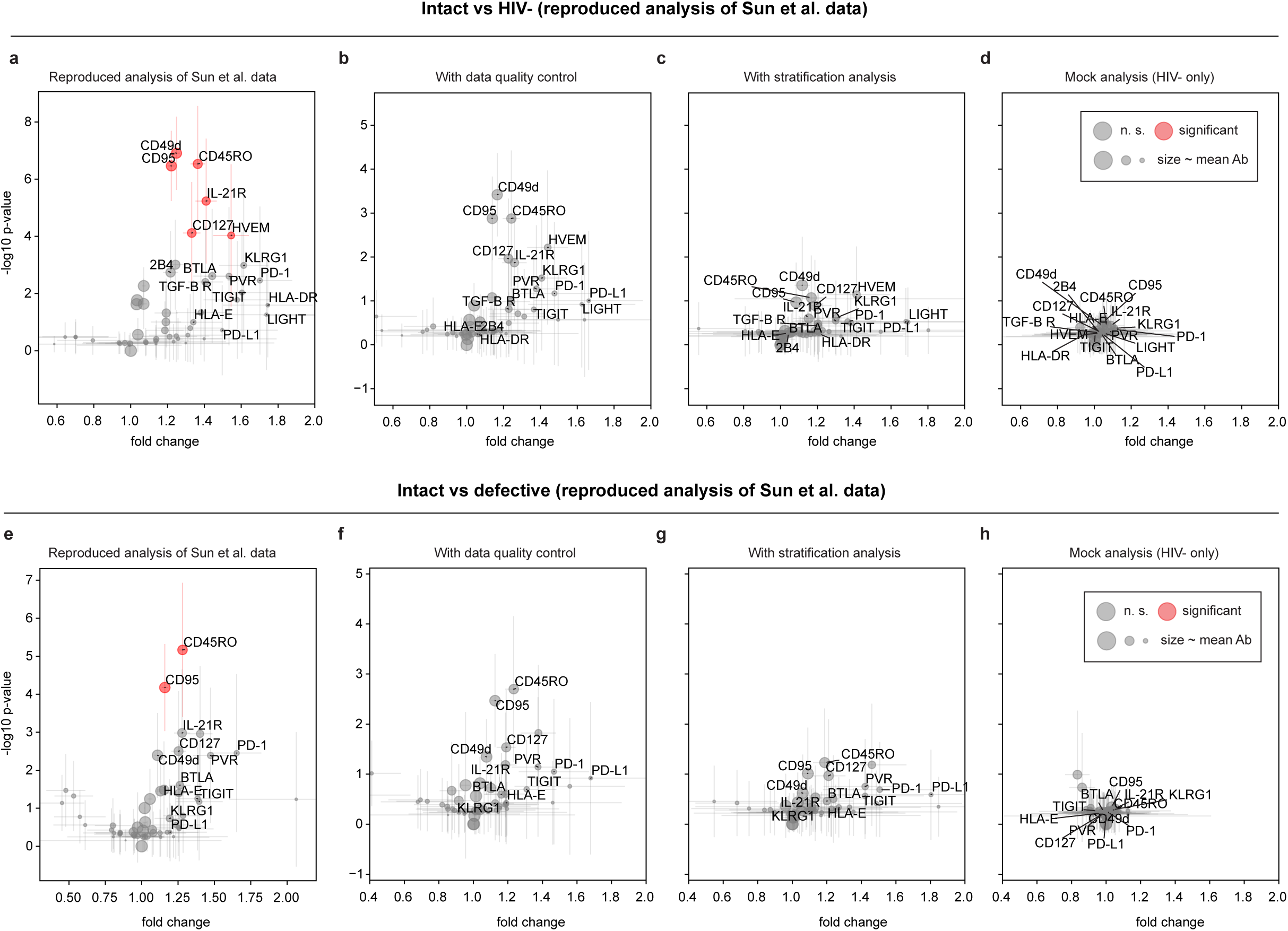
Re-analysis of Sun et al. (2023) differential expression data. **a-d)** Cells with intact provirus versus non-infected cells. **a)** Volcano plot replicating published analysis in Sun et al. (2023) with their data and our implementation of their differential expression analysis method. **b)** Same analysis after the removal of low-quality cells. **c)** Our analysis of Sun et al data using stratification to remove cell type correlations. **d)** Mock analysis of Sun et al data. **e-h)** Cells with intact provirus versus cells with defective virus. **e)** Analogous to **e**, volcano plot replicating the analysis published in Sun et al. (2023) **f)** Same analysis after removal of low-quality cells. **g)** Our analysis of Sun et al data using stratification to remove cell type correlations. **h)** Mock analysis of Sun et al data.

**Supplemental Figure 17.**
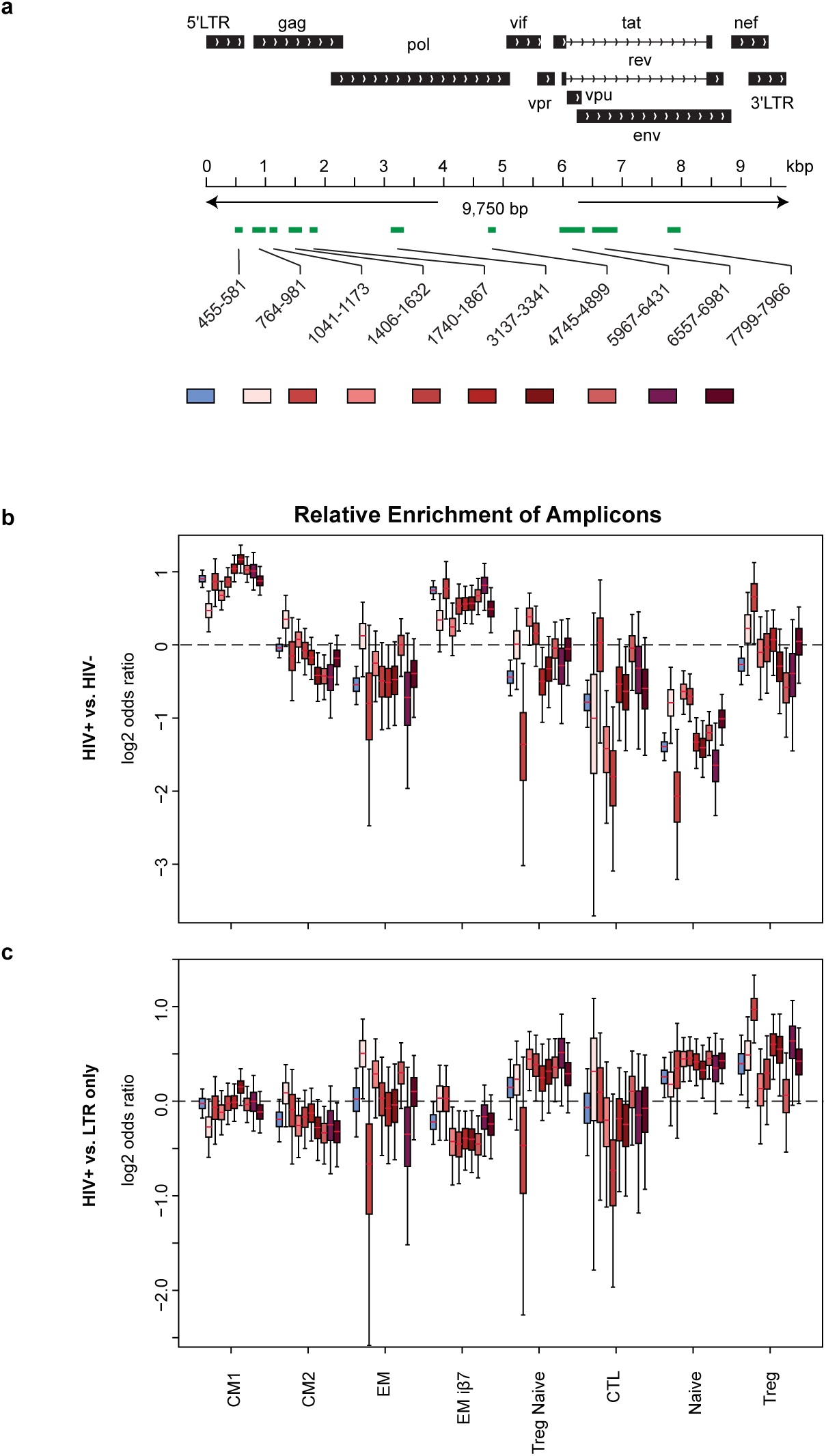
Differences in the Amplicon Landscape across CD4 Subsets. **a)** Location of HIV amplicons across the HXB2 reference genome. **b,c)** Odds ratios of single HIV genome PCR amplicons within CD4 subsets using HIV- (b) or LTR-only (c) cells as the comparator. Most HIV+ cells have many amplicons and thus contribute to multiple groups.

## References

1. Whitney JB, Hill AL, Sanisetty S, Penaloza-MacMaster P, Liu J, Shetty M, Parenteau L, Cabral C, Shields J, Blackmore S, Smith JY, Brinkman AL, Peter LE, Mathew SI, Smith KM, Borducchi EN, Rosenbloom DI, Lewis MG, Hattersley J, Li B, Hesselgesser J, Geleziunas R, Robb ML, Kim JH, Michael NL, Barouch DH. Rapid seeding of the viral reservoir prior to SIV viraemia in rhesus monkeys. Nature. 2014;512(7512):74–7. Epub 20140720. doi: 10.1038/nature13594. PubMed PMID: 25042999; PMCID: PMC4126858.

2. Gantner P, Buranapraditkun S, Pagliuzza A, Dufour C, Pardons M, Mitchell JL, Kroon E, Sacdalan C, Tulmethakaan N, Pinyakorn S, Robb ML, Phanuphak N, Ananworanich J, Hsu D, Vasan S, Trautmann L, Fromentin R, Chomont N. HIV rapidly targets a diverse pool of CD4(+) T cells to establish productive and latent infections. Immunity. 2023;56(3):653–68 e5. Epub 20230217. doi: 10.1016/j.immuni.2023.01.030. PubMed PMID: 36804957; PMCID: PMC10023508.

3. Chaillon A, Gianella S, Dellicour S, Rawlings SA, Schlub TE, De Oliveira MF, Ignacio C, Porrachia M, Vrancken B, Smith DM. HIV persists throughout deep tissues with repopulation from multiple anatomical sources. J Clin Invest. 2020;130(4):1699–712. doi: 10.1172/JCI134815. PubMed PMID: 31910162; PMCID: PMC7108926.

4. Finzi D, Blankson J, Siliciano JD, Margolick JB, Chadwick K, Pierson T, Smith K, Lisziewicz J, Lori F, Flexner C, Quinn TC, Chaisson RE, Rosenberg E, Walker B, Gange S, Gallant J, Siliciano RF. Latent infection of CD4+ T cells provides a mechanism for lifelong persistence of HIV-1, even in patients on effective combination therapy. Nat Med. 1999;5(5):512–7. doi: 10.1038/8394. PubMed PMID: 10229227.

5. Siliciano JD, Kajdas J, Finzi D, Quinn TC, Chadwick K, Margolick JB, Kovacs C, Gange SJ, Siliciano RF. Long-term follow-up studies confirm the stability of the latent reservoir for HIV-1 in resting CD4+ T cells. Nat Med. 2003;9(6):727–8. Epub 20030518. doi: 10.1038/nm880. PubMed PMID: 12754504.

6. McMyn NF, Varriale J, Fray EJ, Zitzmann C, MacLeod H, Lai J, Singhal A, Moskovljevic M, Garcia MA, Lopez BM, Hariharan V, Rhodehouse K, Lynn K, Tebas P, Mounzer K, Montaner LJ, Benko E, Kovacs C, Hoh R, Simonetti FR, Laird GM, Deeks SG, Ribeiro RM, Perelson AS, Siliciano RF, Siliciano JM. The latent reservoir of inducible, infectious HIV-1 does not decrease despite decades of antiretroviral therapy. J Clin Invest. 2023;133(17). Epub 20230901. doi: 10.1172/JCI171554. PubMed PMID: 37463049; PMCID: PMC10471168.

7. Peluso MJ, Bacchetti P, Ritter KD, Beg S, Lai J, Martin JN, Hunt PW, Henrich TJ, Siliciano JD, Siliciano RF, Laird GM, Deeks SG. Differential decay of intact and defective proviral DNA in HIV-1-infected individuals on suppressive antiretroviral therapy. JCI Insight. 2020;5(4). Epub 20200227. doi: 10.1172/jci.insight.132997. PubMed PMID: 32045386; PMCID: PMC7101154.

8. Simonetti FR, White JA, Tumiotto C, Ritter KD, Cai M, Gandhi RT, Deeks SG, Howell BJ, Montaner LJ, Blankson JN, Martin A, Laird GM, Siliciano RF, Mellors JW, Siliciano JD. Intact proviral DNA assay analysis of large cohorts of people with HIV provides a benchmark for the frequency and composition of persistent proviral DNA. Proc Natl Acad Sci U S A. 2020;117(31):18692–700. Epub 20200720. doi: 10.1073/pnas.2006816117. PubMed PMID: 32690683; PMCID: PMC7414172.

9. Bruner KM, Wang Z, Simonetti FR, Bender AM, Kwon KJ, Sengupta S, Fray EJ, Beg SA, Antar AAR, Jenike KM, Bertagnolli LN, Capoferri AA, Kufera JT, Timmons A, Nobles C, Gregg J, Wada N, Ho YC, Zhang H, Margolick JB, Blankson JN, Deeks SG, Bushman FD, Siliciano JD, Laird GM, Siliciano RF. A quantitative approach for measuring the reservoir of latent HIV-1 proviruses. Nature. 2019;566(7742):120–5. Epub 20190130. doi: 10.1038/s41586-019-0898-8. PubMed PMID: 30700913; PMCID: PMC6447073.

10. Bruner KM, Murray AJ, Pollack RA, Soliman MG, Laskey SB, Capoferri AA, Lai J, Strain MC, Lada SM, Hoh R, Ho YC, Richman DD, Deeks SG, Siliciano JD, Siliciano RF. Defective proviruses rapidly accumulate during acute HIV-1 infection. Nat Med. 2016;22(9):1043–9. Epub 20160808. doi: 10.1038/nm.4156. PubMed PMID: 27500724; PMCID: PMC5014606.

11. Martin AR, Bender AM, Hackman J, Kwon KJ, Lynch BA, Bruno D, Martens C, Beg S, Florman SS, Desai N, Segev D, Laird GM, Siliciano JD, Quinn TC, Tobian AAR, Durand CM, Siliciano RF, Redd AD. Similar Frequency and Inducibility of Intact Human Immunodeficiency Virus-1 Proviruses in Blood and Lymph Nodes. J Infect Dis. 2021;224(2):258–68. doi: 10.1093/infdis/jiaa736. PubMed PMID: 33269401; PMCID: PMC8280486.

12. Pardons M, Baxter AE, Massanella M, Pagliuzza A, Fromentin R, Dufour C, Leyre L, Routy JP, Kaufmann DE, Chomont N. Single-cell characterization and quantification of translation-competent viral reservoirs in treated and untreated HIV infection. PLoS Pathog. 2019;15(2):e1007619. Epub 20190227. doi: 10.1371/journal.ppat.1007619. PubMed PMID: 30811499; PMCID: PMC6411230.

13. Liu R, Yeh YJ, Varabyou A, Collora JA, Sherrill-Mix S, Talbot CC, Jr., Mehta S, Albrecht K, Hao H, Zhang H, Pollack RA, Beg SA, Calvi RM, Hu J, Durand CM, Ambinder RF, Hoh R, Deeks SG, Chiarella J, Spudich S, Douek DC, Bushman FD, Pertea M, Ho YC. Single-cell transcriptional landscapes reveal HIV-1-driven aberrant host gene transcription as a potential therapeutic target. Sci Transl Med. 2020;12(543). doi: 10.1126/scitranslmed.aaz0802. PubMed PMID: 32404504; PMCID: PMC7453882.

14. Hosmane NN, Kwon KJ, Bruner KM, Capoferri AA, Beg S, Rosenbloom DI, Keele BF, Ho YC, Siliciano JD, Siliciano RF. Proliferation of latently infected CD4(+) T cells carrying replication-competent HIV-1: Potential role in latent reservoir dynamics. J Exp Med. 2017;214(4):959–72. Epub 2017/03/28. doi: 10.1084/jem.20170193. PubMed PMID: 28341641; PMCID: PMC5379987.

15. Ho YC, Shan L, Hosmane NN, Wang J, Laskey SB, Rosenbloom DI, Lai J, Blankson JN, Siliciano JD, Siliciano RF. Replication-competent noninduced proviruses in the latent reservoir increase barrier to HIV-1 cure. Cell. 2013;155(3):540–51. Epub 20131024. doi: 10.1016/j.cell.2013.09.020. PubMed PMID: 24243014; PMCID: PMC3896327.

16. Einkauf KB, Osborn MR, Gao C, Sun W, Sun X, Lian X, Parsons EM, Gladkov GT, Seiger KW, Blackmer JE, Jiang C, Yukl SA, Rosenberg ES, Yu XG, Lichterfeld M. Parallel analysis of transcription, integration, and sequence of single HIV-1 proviruses. Cell. 2022;185(2):266–82 e15. Epub 20220112. doi: 10.1016/j.cell.2021.12.011. PubMed PMID: 35026153; PMCID: PMC8809251.

17. Wu VH, Nordin JML, Nguyen S, Joy J, Mampe F, Del Rio Estrada PM, Torres-Ruiz F, Gonzalez-Navarro M, Luna-Villalobos YA, Avila-Rios S, Reyes-Teran G, Tebas P, Montaner LJ, Bar KJ, Vella LA, Betts MR. Profound phenotypic and epigenetic heterogeneity of the HIV-1-infected CD4(+) T cell reservoir. Nat Immunol. 2023;24(2):359–70. Epub 20221219. doi: 10.1038/s41590-022-01371-3. PubMed PMID: 36536105; PMCID: PMC9892009.

18. Demaree B, Delley CL, Vasudevan HN, Peretz CAC, Ruff D, Smith CC, Abate AR. Joint profiling of DNA and proteins in single cells to dissect genotype-phenotype associations in leukemia. Nat Commun. 2021;12(1):1583. Epub 20210311. doi: 10.1038/s41467-021-21810-3. PubMed PMID: 33707421; PMCID: PMC7952600.

19. Sun W, Gao C, Hartana CA, Osborn MR, Einkauf KB, Lian X, Bone B, Bonheur N, Chun TW, Rosenberg ES, Walker BD, Yu XG, Lichterfeld M. Phenotypic signatures of immune selection in HIV-1 reservoir cells. Nature. 2023;614(7947):309–17. Epub 20230104. doi: 10.1038/s41586-022-05538-8. PubMed PMID: 36599977; PMCID: PMC9908552.

20. Hiener B, Horsburgh BA, Eden JS, Barton K, Schlub TE, Lee E, von Stockenstrom S, Odevall L, Milush JM, Liegler T, Sinclair E, Hoh R, Boritz EA, Douek D, Fromentin R, Chomont N, Deeks SG, Hecht FM, Palmer S. Identification of Genetically Intact HIV-1 Proviruses in Specific CD4(+) T Cells from Effectively Treated Participants. Cell Rep. 2017;21(3):813–22. doi: 10.1016/j.celrep.2017.09.081. PubMed PMID: 29045846; PMCID: PMC5960642.

21. Delley CL, Abate AR. Modular barcode beads for microfluidic single cell genomics. Sci Rep. 2021;11(1):10857. Epub 20210525. doi: 10.1038/s41598-021-90255-x. PubMed PMID: 34035349; PMCID: PMC8149635.

22. Fleming SJ, Chaffin MD, Arduini A, Akkad AD, Banks E, Marioni JC, Philippakis AA, Ellinor PT, Babadi M. Unsupervised removal of systematic background noise from droplet-based single-cell experiments using CellBender. Nat Methods. 2023;20(9):1323–35. Epub 20230807. doi: 10.1038/s41592-023-01943-7. PubMed PMID: 37550580.

23. Gayoso A, Steier Z, Lopez R, Regier J, Nazor KL, Streets A, Yosef N. Joint probabilistic modeling of single-cell multi-omic data with totalVI. Nat Methods. 2021;18(3):272–82. Epub 20210215. doi: 10.1038/s41592-020-01050-x. PubMed PMID: 33589839; PMCID: PMC7954949.

24. Lopez R, Regier J, Cole MB, Jordan MI, Yosef N. Deep generative modeling for single-cell transcriptomics. Nat Methods. 2018;15(12):1053–8. Epub 20181130. doi: 10.1038/s41592-018-0229-2. PubMed PMID: 30504886; PMCID: PMC6289068.

25. Gayoso A, Lopez R, Xing G, Boyeau P, Valiollah Pour Amiri V, Hong J, Wu K, Jayasuriya M, Mehlman E, Langevin M, Liu Y, Samaran J, Misrachi G, Nazaret A, Clivio O, Xu C, Ashuach T, Gabitto M, Lotfollahi M, Svensson V, da Veiga Beltrame E, Kleshchevnikov V, Talavera-Lopez C, Pachter L, Theis FJ, Streets A, Jordan MI, Regier J, Yosef N. A Python library for probabilistic analysis of single-cell omics data. Nat Biotechnol. 2022;40(2):163–6. doi: 10.1038/s41587-021-01206-w. PubMed PMID: 35132262.

26. Traag VA, Waltman L, van Eck NJ. From Louvain to Leiden: guaranteeing well-connected communities. Sci Rep. 2019;9(1):5233. Epub 20190326. doi: 10.1038/s41598-019-41695-z. PubMed PMID: 30914743; PMCID: PMC6435756.

27. Kotliarov Y, Sparks R, Martins AJ, Mule MP, Lu Y, Goswami M, Kardava L, Banchereau R, Pascual V, Biancotto A, Chen J, Schwartzberg PL, Bansal N, Liu CC, Cheung F, Moir S, Tsang JS. Broad immune activation underlies shared set point signatures for vaccine responsiveness in healthy individuals and disease activity in patients with lupus. Nat Med. 2020;26(4):618–29. Epub 20200224. doi: 10.1038/s41591-020-0769-8. PubMed PMID: 32094927; PMCID: PMC8392163.

28. Hao Y, Hao S, Andersen-Nissen E, Mauck WM, 3rd, Zheng S, Butler A, Lee MJ, Wilk AJ, Darby C, Zager M, Hoffman P, Stoeckius M, Papalexi E, Mimitou EP, Jain J, Srivastava A, Stuart T, Fleming LM, Yeung B, Rogers AJ, McElrath JM, Blish CA, Gottardo R, Smibert P, Satija R. Integrated analysis of multimodal single-cell data. Cell. 2021;184(13):3573–87 e29. Epub 20210531. doi: 10.1016/j.cell.2021.04.048. PubMed PMID: 34062119; PMCID: PMC8238499.

29. Anderson EM, Simonetti FR, Gorelick RJ, Hill S, Gouzoulis MA, Bell J, Rehm C, Perez L, Boritz E, Wu X, Wells D, Hughes SH, Rao V, Coffin JM, Kearney MF, Maldarelli F. Dynamic Shifts in the HIV Proviral Landscape During Long Term Combination Antiretroviral Therapy: Implications for Persistence and Control of HIV Infections. Viruses. 2020;12(2). Epub 20200125. doi: 10.3390/v12020136. PubMed PMID: 31991737; PMCID: PMC7077288.

30. Botha JC, Demirov D, Gordijn C, Katusiime MG, Bale MJ, Wu X, Wells D, Hughes SH, Cotton MF, Mellors JW, Kearney MF, van Zyl GU. The largest HIV-1-infected T cell clones in children on long-term combination antiretroviral therapy contain solo LTRs. mBio. 2023;14(4):e0111623. Epub 20230802. doi: 10.1128/mbio.01116-23. PubMed PMID: 37530525; PMCID: PMC10470503.

31. Arthos J, Cicala C, Nawaz F, Byrareddy SN, Villinger F, Santangelo PJ, Ansari AA, Fauci AS. The Role of Integrin alpha(4)beta(7) in HIV Pathogenesis and Treatment. Curr HIV/AIDS Rep. 2018;15(2):127–35. doi: 10.1007/s11904-018-0382-3. PubMed PMID: 29478152; PMCID: PMC5882766.

32. Li X, Liu Z, Li Q, Hu R, Zhao L, Yang Y, Zhao J, Huang Z, Gao H, Li L, Cai W, Deng K. CD161(+) CD4(+) T Cells Harbor Clonally Expanded Replication-Competent HIV-1 in Antiretroviral Therapy-Suppressed Individuals. mBio. 2019;10(5). Epub 20191008. doi: 10.1128/mBio.02121-19. PubMed PMID: 31594817; PMCID: PMC6786872.

33. Wei Y, Davenport TC, Collora JA, Ma HK, Pinto-Santini D, Lama J, Alfaro R, Duerr A, Ho YC. Single-cell epigenetic, transcriptional, and protein profiling of latent and active HIV-1 reservoir revealed that IKZF3 promotes HIV-1 persistence. Immunity. 2023;56(11):2584–601 e7. Epub 20231102. doi: 10.1016/j.immuni.2023.10.002. PubMed PMID: 37922905; PMCID: PMC10843106.

34. Gosselin A, Wiche Salinas TR, Planas D, Wacleche VS, Zhang Y, Fromentin R, Chomont N, Cohen EA, Shacklett B, Mehraj V, Ghali MP, Routy JP, Ancuta P. HIV persists in CCR6+CD4+ T cells from colon and blood during antiretroviral therapy. AIDS. 2017;31(1):35–48. doi: 10.1097/QAD.0000000000001309. PubMed PMID: 27835617; PMCID: PMC5131694.

35. Anderson JL, Khoury G, Fromentin R, Solomon A, Chomont N, Sinclair E, Milush JM, Hartogensis W, Bacchetti P, Roche M, Tumpach C, Gartner M, Pitman MC, Epling CL, Hoh R, Hecht FM, Somsouk M, Cameron PU, Deeks SG, Lewin SR. Human Immunodeficiency Virus (HIV)-Infected CCR6+ Rectal CD4+ T Cells and HIV Persistence On Antiretroviral Therapy. J Infect Dis. 2020;221(5):744–55. doi: 10.1093/infdis/jiz509. PubMed PMID: 31796951; PMCID: PMC7026892.

36. Maggi L, Santarlasci V, Capone M, Peired A, Frosali F, Crome SQ, Querci V, Fambrini M, Liotta F, Levings MK, Maggi E, Cosmi L, Romagnani S, Annunziato F. CD161 is a marker of all human IL-17-producing T-cell subsets and is induced by RORC. Eur J Immunol. 2010;40(8):2174–81. doi: 10.1002/eji.200940257. PubMed PMID: 20486123.

37. Dufour C, Richard C, Pardons M, Massanella M, Ackaoui A, Murrell B, Routy B, Thomas R, Routy JP, Fromentin R, Chomont N. Phenotypic characterization of single CD4+ T cells harboring genetically intact and inducible HIV genomes. Nat Commun. 2023;14(1):1115. Epub 20230227. doi: 10.1038/s41467-023-36772-x. PubMed PMID: 36849523; PMCID: PMC9971253.

38. Nawaz F, Cicala C, Van Ryk D, Block KE, Jelicic K, McNally JP, Ogundare O, Pascuccio M, Patel N, Wei D, Fauci AS, Arthos J. The genotype of early-transmitting HIV gp120s promotes alpha (4) beta(7)-reactivity, revealing alpha (4) beta(7) +/CD4+ T cells as key targets in mucosal transmission. PLoS Pathog. 2011;7(2):e1001301. Epub 20110224. doi: 10.1371/journal.ppat.1001301. PubMed PMID: 21383973; PMCID: PMC3044691.

39. Veazey RS, Marx PA, Lackner AA. The mucosal immune system: primary target for HIV infection and AIDS. Trends Immunol. 2001;22(11):626–33. doi: 10.1016/s1471-4906(01)02039-7. PubMed PMID: 11698224.

40. Tokarev A, McKinnon LR, Pagliuzza A, Sivro A, Omole TE, Kroon E, Chomchey N, Phanuphak N, Schuetz A, Robb ML, Eller MA, Ananworanich J, Chomont N, Bolton DL. Preferential Infection of alpha4beta7+ Memory CD4+ T Cells During Early Acute Human Immunodeficiency Virus Type 1 Infection. Clin Infect Dis. 2020;71(11):e735–e43. doi: 10.1093/cid/ciaa497. PubMed PMID: 32348459; PMCID: PMC7778353.

41. Sivro A, Schuetz A, Sheward D, Joag V, Yegorov S, Liebenberg LJ, Yende-Zuma N, Stalker A, Mwatelah RS, Selhorst P, Garrett N, Samsunder N, Balgobin A, Nawaz F, Cicala C, Arthos J, Fauci AS, Anzala AO, Kimani J, Bagaya BS, Kiwanuka N, Williamson C, Kaul R, Passmore JS, Phanuphak N, Ananworanich J, Ansari A, Abdool Karim Q, Abdool Karim SS, McKinnon LR, Caprisa, groups RVs. Integrin alpha(4)beta(7) expression on peripheral blood CD4(+) T cells predicts HIV acquisition and disease progression outcomes. Sci Transl Med. 2018;10(425). doi: 10.1126/scitranslmed.aam6354. PubMed PMID: 29367348; PMCID: PMC6820005.

42. Parrish NF, Wilen CB, Banks LB, Iyer SS, Pfaff JM, Salazar-Gonzalez JF, Salazar MG, Decker JM, Parrish EH, Berg A, Hopper J, Hora B, Kumar A, Mahlokozera T, Yuan S, Coleman C, Vermeulen M, Ding H, Ochsenbauer C, Tilton JC, Permar SR, Kappes JC, Betts MR, Busch MP, Gao F, Montefiori D, Haynes BF, Shaw GM, Hahn BH, Doms RW. Transmitted/founder and chronic subtype C HIV-1 use CD4 and CCR5 receptors with equal efficiency and are not inhibited by blocking the integrin alpha4beta7. PLoS Pathog. 2012;8(5):e1002686. Epub 20120531. doi: 10.1371/journal.ppat.1002686. PubMed PMID: 22693444; PMCID: PMC3364951.

43. Card CM, Abrenica B, McKinnon LR, Ball TB, Su RC. Endothelial Cells Promote Productive HIV Infection of Resting CD4(+) T Cells by an Integrin-Mediated Cell Adhesion-Dependent Mechanism. AIDS Res Hum Retroviruses. 2022;38(2):111–26. Epub 20211011. doi: 10.1089/AID.2021.0034. PubMed PMID: 34465136; PMCID: PMC8861939.

44. Martinelli E, Veglia F, Goode D, Guerra-Perez N, Aravantinou M, Arthos J, Piatak M, Jr., Lifson JD, Blanchard J, Gettie A, Robbiani M. The frequency of alpha(4)beta(7)(high) memory CD4(+) T cells correlates with susceptibility to rectal simian immunodeficiency virus infection. J Acquir Immune Defic Syndr. 2013;64(4):325–31. doi: 10.1097/QAI.0b013e31829f6e1a. PubMed PMID: 23797688; PMCID: PMC3815485.

45. Bacchus-Souffan C, Fitch M, Symons J, Abdel-Mohsen M, Reeves DB, Hoh R, Stone M, Hiatt J, Kim P, Chopra A, Ahn H, York VA, Cameron DL, Hecht FM, Martin JN, Yukl SA, Mallal S, Cameron PU, Deeks SG, Schiffer JT, Lewin SR, Hellerstein MK, McCune JM, Hunt PW. Relationship between CD4 T cell turnover, cellular differentiation and HIV persistence during ART. PLoS Pathog. 2021;17(1):e1009214. Epub 20210119. doi: 10.1371/journal.ppat.1009214. PubMed PMID: 33465157; PMCID: PMC7846027.

46. Fromentin R, Bakeman W, Lawani MB, Khoury G, Hartogensis W, DaFonseca S, Killian M, Epling L, Hoh R, Sinclair E, Hecht FM, Bacchetti P, Deeks SG, Lewin SR, Sekaly RP, Chomont N. CD4+ T Cells Expressing PD-1, TIGIT and LAG-3 Contribute to HIV Persistence during ART. PLoS Pathog. 2016;12(7):e1005761. Epub 20160714. doi: 10.1371/journal.ppat.1005761. PubMed PMID: 27415008; PMCID: PMC4944956.

47. Evans VA, van der Sluis RM, Solomon A, Dantanarayana A, McNeil C, Garsia R, Palmer S, Fromentin R, Chomont N, Sekaly RP, Cameron PU, Lewin SR. Programmed cell death-1 contributes to the establishment and maintenance of HIV-1 latency. AIDS. 2018;32(11):1491–7. doi: 10.1097/QAD.0000000000001849. PubMed PMID: 29746296; PMCID: PMC6026054.

48. Wang X, Xu H, Alvarez X, Pahar B, Moroney-Rasmussen T, Lackner AA, Veazey RS. Distinct expression patterns of CD69 in mucosal and systemic lymphoid tissues in primary SIV infection of rhesus macaques. PLoS One. 2011;6(11):e27207. Epub 20111109. doi: 10.1371/journal.pone.0027207. PubMed PMID: 22096538; PMCID: PMC3212564.

49. McKinnon LR, Nyanga B, Chege D, Izulla P, Kimani M, Huibner S, Gelmon L, Block KE, Cicala C, Anzala AO, Arthos J, Kimani J, Kaul R. Characterization of a human cervical CD4+ T cell subset coexpressing multiple markers of HIV susceptibility. J Immunol. 2011;187(11):6032–42. Epub 20111102. doi: 10.4049/jimmunol.1101836. PubMed PMID: 22048765.

50. Iglesias-Ussel M, Vandergeeten C, Marchionni L, Chomont N, Romerio F. High levels of CD2 expression identify HIV-1 latently infected resting memory CD4+ T cells in virally suppressed subjects. J Virol. 2013;87(16):9148–58. Epub 20130612. doi: 10.1128/JVI.01297-13. PubMed PMID: 23760244; PMCID: PMC3754042.

51. Neidleman J, Luo X, Frouard J, Xie G, Hsiao F, Ma T, Morcilla V, Lee A, Telwatte S, Thomas R, Tamaki W, Wheeler B, Hoh R, Somsouk M, Vohra P, Milush J, James KS, Archin NM, Hunt PW, Deeks SG, Yukl SA, Palmer S, Greene WC, Roan NR. Phenotypic analysis of the unstimulated in vivo HIV CD4 T cell reservoir. Elife. 2020;9. Epub 20200929. doi: 10.7554/eLife.60933. PubMed PMID: 32990219; PMCID: PMC7524554.

52. Sperber HS, Raymond KA, Bouzidi MS, Ma TC, Valdebenito S, Eugenin EA, Roan NR, Deeks SG, Winning S, Fandrey J, Schwarzer R, Pillai SK. The hypoxia-regulated ectonucleotidase CD73 is a host determinant of HIV latency. Cell Rep. 2023;42(11). doi: ARTN 113285 10.1016/j.celrep.2023.113285. PubMed PMID: WOS:001105714700001.

53. Dahmane S, Doucet C, Le Gall A, Chamontin C, Dosset P, Murcy F, Fernandez L, Salas D, Rubinstein E, Mougel M, Nollmann M, Milhiet PE. Nanoscale organization of tetraspanins during HIV-1 budding by correlative dSTORM/AFM. Nanoscale. 2019;11(13):6036–44. doi: 10.1039/c8nr07269h. PubMed PMID: WOS:000464518400023.

54. Sims B, Farrow AL, Williams SD, Bansal A, Krendelchtchikov A, Matthews QL. Tetraspanin blockage reduces exosome-mediated HIV-1 entry. Arch Virol. 2018;163(6):1683–9. doi: 10.1007/s00705-018-3737-6. PubMed PMID: WOS:000432599500031.

55. Florin L, Lang T. Tetraspanin Assemblies in Virus Infection. Front Immunol. 2018;9:1140. Epub 20180525. doi: 10.3389/fimmu.2018.01140. PubMed PMID: 29887866; PMCID: PMC5981178.

56. Shin SW, Mudvari P, Thaploo S, Wheeler MA, Douek DC, Quintana FJ, Boritz EA, Abate AR, Clark IC. FIND-seq: high-throughput nucleic acid cytometry for rare single-cell transcriptomics. Nat Protoc. 2024;19(11):3191–218. doi: 10.1038/s41596-024-01021-y. PubMed PMID: WOS:001273949100001.

57. Clark IC, Mudvari P, Thaploo S, Smith S, Abu-Laban M, Hamouda M, Theberge M, Shah SK, Ko SH, Pérez L, Bunis DG, Lee JS, Kilam D, Zakaria S, Choi S, Darko S, Henry AR, Wheeler MA, Hoh R, Butrus S, Deeks SG, Quintana FJ, Douek DC, Abate AR, Boritz EA. HIV silencing and cell survival signatures in infected T cell reservoirs. Nature. 2023;614(7947). doi: 10.1038/s41586-022-05556-6. PubMed PMID: WOS:000952315900008.

58. Einkauf KB, Lee GQ, Gao C, Sharaf R, Sun X, Hua S, Chen SM, Jiang C, Lian X, Chowdhury FZ, Rosenberg ES, Chun TW, Li JZ, Yu XG, Lichterfeld M. Intact HIV-1 proviruses accumulate at distinct chromosomal positions during prolonged antiretroviral therapy. J Clin Invest. 2019;129(3):988–98. Epub 20190128. doi: 10.1172/JCI124291. PubMed PMID: 30688658; PMCID: PMC6391088.

59. Zerbato JM, McMahon DK, Sobolewski MD, Mellors JW, Sluis-Cremer N. Naive CD4+ T Cells Harbor a Large Inducible Reservoir of Latent, Replication-competent Human Immunodeficiency Virus Type 1. Clin Infect Dis. 2019;69(11):1919–25. doi: 10.1093/cid/ciz108. PubMed PMID: 30753360; PMCID: PMC6853701.

60. Venanzi Rullo E, Pinzone MR, Cannon L, Weissman S, Ceccarelli M, Zurakowski R, Nunnari G, O’Doherty U. Persistence of an intact HIV reservoir in phenotypically naive T cells. JCI Insight. 2020;5(20). Epub 20201015. doi: 10.1172/jci.insight.133157. PubMed PMID: 33055422; PMCID: PMC7605525.

61. Mavigner M, Habib J, Deleage C, Rosen E, Mattingly C, Bricker K, Kashuba A, Amblard F, Schinazi RF, Lawson B, Vanderford TH, Jean S, Cohen J, McGary C, Paiardini M, Wood MP, Sodora DL, Silvestri G, Estes J, Chahroudi A. Simian Immunodeficiency Virus Persistence in Cellular and Anatomic Reservoirs in Antiretroviral Therapy-Suppressed Infant Rhesus Macaques. J Virol. 2018;92(18). Epub 20180829. doi: 10.1128/JVI.00562-18. PubMed PMID: 29997216; PMCID: PMC6146711.

62. Obregon-Perko V, Bricker KM, Mensah G, Uddin F, Kumar MR, Fray EJ, Siliciano RF, Schoof N, Horner A, Mavigner M, Liang S, Vanderford T, Sass J, Chan C, Berendam SJ, Bar KJ, Shaw GM, Silvestri G, Fouda GG, Permar SR, Chahroudi A. Simian-Human Immunodeficiency Virus SHIV.C.CH505 Persistence in ART-Suppressed Infant Macaques Is Characterized by Elevated SHIV RNA in the Gut and a High Abundance of Intact SHIV DNA in Naive CD4(+) T Cells. J Virol. 2020;95(2). Epub 20201222. doi: 10.1128/JVI.01669-20. PubMed PMID: 33087463; PMCID: PMC7944446.

63. Rullo EV, Cannon L, Pinzone MR, Ceccarelli M, Nunnari G, O’Doherty U. Genetic Evidence That Naive T Cells Can Contribute Significantly to the Human Immunodeficiency Virus Intact Reservoir: Time to Re-evaluate Their Role. Clinical Infectious Diseases. 2019;69(12):2236–7. doi: 10.1093/cid/ciz378. PubMed PMID: WOS:000501729300043.

64. Reeves DB, Bacchus-Souffan C, Fitch M, Abdel-Mohsen M, Hoh R, Ahn H, Stone M, Hecht F, Martin J, Deeks SG, Hellerstein MK, McCune JM, Schiffer JT, Hunt PW. Estimating the contribution of CD4 T cell subset proliferation and differentiation to HIV persistence. Nat Commun. 2023;14(1):6145. Epub 20231002. doi: 10.1038/s41467-023-41521-1. PubMed PMID: 37783718; PMCID: PMC10545742.

65. Potapov V, Ong JL, Kucera RB, Langhorst BW, Bilotti K, Pryor JM, Cantor EJ, Canton B, Knight TF, Evans TC, Jr., Lohman GJS. Comprehensive Profiling of Four Base Overhang Ligation Fidelity by T4 DNA Ligase and Application to DNA Assembly. ACS Synth Biol. 2018;7(11):2665–74. Epub 20181029. doi: 10.1021/acssynbio.8b00333. PubMed PMID: 30335370.

66. Hug H, Schuler R. Measurement of the number of molecules of a single mRNA species in a complex mRNA preparation. J Theor Biol. 2003;221(4):615–24. doi: 10.1006/jtbi.2003.3211. PubMed PMID: 12713944.

67. Shalek AK, Satija R, Adiconis X, Gertner RS, Gaublomme JT, Raychowdhury R, Schwartz S, Yosef N, Malboeuf C, Lu D, Trombetta JJ, Gennert D, Gnirke A, Goren A, Hacohen N, Levin JZ, Park H, Regev A. Single-cell transcriptomics reveals bimodality in expression and splicing in immune cells. Nature. 2013;498(7453):236–40. Epub 20130519. doi: 10.1038/nature12172. PubMed PMID: 23685454; PMCID: PMC3683364.

68. Levenshtein VI. Binary codes capable of correcting deletions, insertions, and reversals. [New York]: American Institute of Physics.; 1966. p. 707–10.

69. Melsted P, Ntranos V, Pachter L. The barcode, UMI, set format and BUStools. Bioinformatics. 2019;35(21):4472–3. doi: 10.1093/bioinformatics/btz279. PubMed PMID: 31073610.

70. Melsted P, Booeshaghi AS, Liu L, Gao F, Lu L, Min KHJ, da Veiga Beltrame E, Hjorleifsson KE, Gehring J, Pachter L. Modular, efficient and constant-memory single-cell RNA-seq preprocessing. Nat Biotechnol. 2021;39(7):813–8. Epub 20210401. doi: 10.1038/s41587-021-00870-2. PubMed PMID: 33795888.

71. Smith T, Heger A, Sudbery I. UMI-tools: modeling sequencing errors in Unique Molecular Identifiers to improve quantification accuracy. Genome Res. 2017;27(3):491–9. Epub 20170118. doi: 10.1101/gr.209601.116. PubMed PMID: 28100584; PMCID: PMC5340976.

72. Stoeckius M, Hafemeister C, Stephenson W, Houck-Loomis B, Chattopadhyay PK, Swerdlow H, Satija R, Smibert P. Simultaneous epitope and transcriptome measurement in single cells. Nat Methods. 2017;14(9):865–8. Epub 20170731. doi: 10.1038/nmeth.4380. PubMed PMID: 28759029; PMCID: PMC5669064.

73. Wolf FA, Angerer P, Theis FJ. SCANPY: large-scale single-cell gene expression data analysis. Genome Biol. 2018;19(1):15. Epub 20180206. doi: 10.1186/s13059-017-1382-0. PubMed PMID: 29409532; PMCID: PMC5802054.

74. McInnes LaH, J. UMAP: uniform manifold approximation and projection for dimension reduction. https://arxivorg/abs/180203426 2020.

75. Xu C, Lopez R, Mehlman E, Regier J, Jordan MI, Yosef N. Probabilistic harmonization and annotation of single-cell transcriptomics data with deep generative models. Mol Syst Biol. 2021;17(1):e9620. doi: 10.15252/msb.20209620. PubMed PMID: 33491336; PMCID: PMC7829634.

76. Tabula Sapiens C, Jones RC, Karkanias J, Krasnow MA, Pisco AO, Quake SR, Salzman J, Yosef N, Bulthaup B, Brown P, Harper W, Hemenez M, Ponnusamy R, Salehi A, Sanagavarapu BA, Spallino E, Aaron KA, Concepcion W, Gardner JM, Kelly B, Neidlinger N, Wang Z, Crasta S, Kolluru S, Morri M, Tan SY, Travaglini KJ, Xu C, Alcantara-Hernandez M, Almanzar N, Antony J, Beyersdorf B, Burhan D, Calcuttawala K, Carter MM, Chan CKF, Chang CA, Chang S, Colville A, Culver RN, Cvijovic I, D’Amato G, Ezran C, Galdos FX, Gillich A, Goodyer WR, Hang Y, Hayashi A, Houshdaran S, Huang X, Irwin JC, Jang S, Juanico JV, Kershner AM, Kim S, Kiss B, Kong W, Kumar ME, Kuo AH, Li B, Loeb GB, Lu WJ, Mantri S, Markovic M, McAlpine PL, de Morree A, Mrouj K, Mukherjee S, Muser T, Neuhofer P, Nguyen TD, Perez K, Puluca N, Qi Z, Rao P, Raquer-McKay H, Schaum N, Scott B, Seddighzadeh B, Segal J, Sen S, Sikandar S, Spencer SP, Steffes LC, Subramaniam VR, Swarup A, Swift M, Van Treuren W, Trimm E, Veizades S, Vijayakumar S, Vo KC, Vorperian SK, Wang W, Weinstein HNW, Winkler J, Wu TTH, Xie J, Yung AR, Zhang Y, Detweiler AM, Mekonen H, Neff NF, Sit RV, Tan M, Yan J, Bean GR, Charu V, Forgo E, Martin BA, Ozawa MG, Silva O, Toland A, Vemuri VNP, Afik S, Awayan K, Botvinnik OB, Byrne A, Chen M, Dehghannasiri R, Gayoso A, Granados AA, Li Q, Mahmoudabadi G, McGeever A, Olivieri JE, Park M, Ravikumar N, Stanley G, Tan W, Tarashansky AJ, Vanheusden R, Wang P, Wang S, Xing G, Dethlefsen L, Ezran C, Gillich A, Hang Y, Ho PY, Irwin JC, Jang S, Leylek R, Liu S, Maltzman JS, Metzger RJ, Phansalkar R, Sasagawa K, Sinha R, Song H, Swarup A, Trimm E, Veizades S, Wang B, Beachy PA, Clarke MF, Giudice LC, Huang FW, Huang KC, Idoyaga J, Kim SK, Kuo CS, Nguyen P, Rando TA, Red-Horse K, Reiter J, Relman DA, Sonnenburg JL, Wu A, Wu SM, Wyss-Coray T. The Tabula Sapiens: A multiple-organ, single-cell transcriptomic atlas of humans. Science. 2022;376(6594):eabl4896. Epub 20220513. doi: 10.1126/science.abl4896. PubMed PMID: 35549404; PMCID: PMC9812260.

77. Langmead B, Salzberg SL. Fast gapped-read alignment with Bowtie 2. Nat Methods. 2012;9(4):357–9. Epub 20120304. doi: 10.1038/nmeth.1923. PubMed PMID: 22388286; PMCID: PMC3322381.

78. Macosko EZ, Basu A, Satija R, Nemesh J, Shekhar K, Goldman M, Tirosh I, Bialas AR, Kamitaki N, Martersteck EM, Trombetta JJ, Weitz DA, Sanes JR, Shalek AK, Regev A, McCarroll SA. Highly Parallel Genome-wide Expression Profiling of Individual Cells Using Nanoliter Droplets. Cell. 2015;161(5):1202–14. doi: 10.1016/j.cell.2015.05.002. PubMed PMID: 26000488; PMCID: PMC4481139.

79. Van der Auwera GA, Carneiro MO, Hartl C, Poplin R, Del Angel G, Levy-Moonshine A, Jordan T, Shakir K, Roazen D, Thibault J, Banks E, Garimella KV, Altshuler D, Gabriel S, DePristo MA. From FastQ data to high confidence variant calls: the Genome Analysis Toolkit best practices pipeline. Curr Protoc Bioinformatics. 2013;43(1110):11 0 1– 0 33. doi: 10.1002/0471250953.bi1110s43. PubMed PMID: 25431634; PMCID: PMC4243306.

80. Love MI, Huber W, Anders S. Moderated estimation of fold change and dispersion for RNA-seq data with DESeq2. Genome Biol. 2014;15(12):550. doi: 10.1186/s13059-014-0550-8. PubMed PMID: 25516281; PMCID: PMC4302049.

81. Robinson MD, McCarthy DJ, Smyth GK. edgeR: a Bioconductor package for differential expression analysis of digital gene expression data. Bioinformatics. 2010;26(1):139–40. Epub 20091111. doi: 10.1093/bioinformatics/btp616. PubMed PMID: 19910308; PMCID: PMC2796818.

82. Hoffman MD, and Andrew Gelman. The No-U-Turn Sampler: Adaptively Setting Path Lengths in Hamiltonian Monte Carlo. arXiv:11114246 [statCO]. 2011.

83. Gelman A, Carlin J. Beyond Power Calculations: Assessing Type S (Sign) and Type M (Magnitude) Errors. Perspect Psychol Sci. 2014;9(6):641–51. doi: 10.1177/1745691614551642. PubMed PMID: 26186114.

